# Constitutive Steroidal Glycoalkaloid Biosynthesis in Tomato is Regulated by the Clade IIIe Basic Helix-Loop-Helix Transcription Factors MYC1 and MYC2

**DOI:** 10.1101/2020.01.27.921833

**Authors:** Gwen Swinnen, Margaux De Meyer, Jacob Pollier, Francisco Javier Molina-Hidalgo, Evi Ceulemans, Rebecca De Clercq, Robin Vanden Bossche, Patricia Fernández-Calvo, Mily Ron, Laurens Pauwels, Alain Goossens

**Author notes:** Corresponding author: Alain Goossens, VIB-UGent Center for Plant Systems Biology, Technologiepark 71, B-9052 Gent (Belgium). Centre for Plant Biotechnology and Genomics, Parque Cientifico y Tecnologico, UPM Campus de Montegancedo, 28223 Madrid, Spain. The author(s) responsible for distribution of materials integral to the findings presented in this article in accordance with the policy described in the Instructions for Authors (www.plantcell.org) is (are): Alain Goossens. Online version contains Web-only data.

## Abstract

Specialized metabolites are produced by plants to fend off biotic enemies. Across the plant kingdom, the biosynthesis of these defense compounds is promoted by jasmonate signaling in which clade IIIe basic helix-loop-helix (bHLH) transcription factors take on a central role. Tomato (*Solanum lycopersicum*) produces cholesterol-derived steroidal glycoalkaloids (SGAs) that act as phytoanticipins against a broad variety of herbivores and pathogens. The biosynthesis of SGAs from cholesterol occurs constitutively in tomato plants and can be further stimulated by jasmonates. Here, we demonstrate that the two tomato clade IIIe bHLH transcription factors, MYC1 and MYC2, redundantly and specifically control the constitutive biosynthesis of SGAs. Double *myc1 myc2* loss-of-function tomato hairy roots displayed suppressed constitutive expression of cholesterol and SGA biosynthesis genes, and consequently severely reduced levels of the main tomato SGAs α-tomatine and dehydrotomatine. In contrast, basal expression of genes involved in canonical jasmonate signaling or in the biosynthesis of highly jasmonate-inducible phenylpropanoid-polyamine conjugates was not affected. Furthermore, CRISPR-Cas9(VQR)-mediated genome editing of a specific *cis*-regulatory element, targeted by MYC1/2, in the promoter of a cholesterol biosynthesis gene led to decreased constitutive expression of this gene, but did not affect its jasmonate inducibility. Our results demonstrate that clade IIIe bHLH transcriptional regulators might have evolved to regulate the biosynthesis of specific constitutively accumulating specialized metabolites independent of jasmonate signaling.

**One sentence summary:** The clade IIIe basic helix-loop-helix transcription factors MYC1 and MYC2 control the constitutive biosynthesis of tomato steroidal glycoalkaloids and might do so independently of jasmonate signaling.

## INTRODUCTION

Plants produce species-specific specialized metabolites to defend themselves against natural enemies, such as herbivores and pathogens. Several *Solanaceae* species, including tomato (*Solanum lycopersicum*), potato (*S. tuberosum*) and eggplant (*S. melongena*), synthesize steroidal glycoalkaloids (SGAs). These nitrogen-containing compounds with a terpenoid skeleton accumulate constitutively in many plant organs, thereby forming a chemical barrier that helps the plant to protect itself against a broad range of biotic agents (Friedman, 2002). The main tomato SGAs, α-tomatine and dehydrotomatine, are present in nearly all plant organs, including leaves, roots, flowers and immature green fruits (Friedman and Levin, 1998; Kozukue et al., 2004). As the fruits mature and ripen, *de novo* SGA synthesis terminates and α-tomatine and dehydrotomatine are converted into esculeosides and dehydroesculeosides, respectively, the predominant SGAs in ripe fruits (Yamanaka et al., 2009; Cárdenas et al., 2019; Nakayasu et al., 2020).

Cholesterol, which in plants is derived from cycloartenol, is the biosynthetic precursor for SGAs. Cycloartenol is produced via the cytosolic mevalonate pathway and forms the branch point between cholesterol and phytosterol biosynthesis. Gene duplication and divergence from phytosterol biosynthetic genes accounts for half of the cholesterogenesis genes (Sawai et al., 2014; Sonawane et al., 2016). The other cholesterol biosynthesis genes are shared between the cholesterol and phytosterol pathway (Sonawane et al., 2016). A set of *GLYCOALKALOID METABOLISM* (*GAME*) genes that are partially organized in metabolic gene clusters, are responsible for the biosynthesis of SGAs from cholesterol (Itkin et al., 2011; Itkin et al., 2013; Sonawane et al., 2018). In a first series of reactions catalyzed by a subset of the GAME proteins, cholesterol is converted into SGA aglycones via multiple biosynthetic steps, including the transamination of the sterol backbone (Itkin et al., 2013; Sonawane et al., 2018). Subsequently, these steroidal alkaloids are glycosylated by uridine diphosphate (UDP) glycosyltransferases to form SGAs (Itkin et al., 2011; Itkin et al., 2013).

The conserved jasmonate (JA) signaling pathway is widely recognized to induce specialized metabolism upon herbivore or necrotrophic pathogen attack in many plant species (De Geyter et al., 2012; Wasternack and Strnad, 2019). The oxylipin-derived JA hormone regulates the expression of its target genes by controlling the activity of certain transcription factors (TFs) that belong to, among others, the basic helix-loop-helix (bHLH) family (Goossens et al., 2017). At low intracellular concentrations of the bioactive (+)-7-*iso*-jasmonoyl-L-isoleucine (JA-Ile) (Fonseca et al., 2009), the activity of these TFs is repressed by the JA ZIM DOMAIN (JAZ) proteins (Chini et al., 2007; Thines et al., 2007; Chini et al., 2016). Elevated intracellular JA-Ile concentrations promote the formation of a co-receptor complex between the JAZ proteins and the F-box protein CORONATINE INSENSITIVE1 (COI1) (Yan et al., 2009; Sheard et al., 2010; Yan et al., 2018), which leads to the ubiquitination and subsequent proteasomal degradation of the interacting JAZ protein (Chini et al., 2007; Thines et al., 2007). This releases the TFs from repression and is followed by the concerted upregulation of multiple genes involved in the same species-specific specialized metabolic pathway (De Geyter et al., 2012; Wasternack and Strnad, 2019).

In tomato and potato, the JA-regulated TF GAME9, which belongs to the APETALA2/Ethylene Response Factor (AP2/ERF) family and is also known as JA-RESPONSIVE ERF 4 (JRE4), is involved in the co-regulation of SGA and cholesterol biosynthesis (Cárdenas et al., 2016; Thagun et al., 2016; Nakayasu et al., 2018). At least for a subset of the cholesterol and SGA biosynthesis genes, GAME9 appears to control their expression in cooperation with tomato MYC2, an ortholog of *Arabidopsis thaliana* (Arabidopsis) MYC2, with GAME9 and MYC2 recognizing GC-rich and G-box elements, respectively, in their target promoters (Cárdenas et al., 2016; Thagun et al., 2016). Arabidopsis MYC2 and its orthologs in other species, including tomato MYC2, are members of the clade IIIe bHLH TFs, which take on a central role in the JA signaling cascade across the plant kingdom (Boter et al., 2004; Kazan and Manners, 2013; Du et al., 2017; Goossens et al., 2017). Besides MYC2, tomato possesses a second clade IIIe bHLH TF, MYC1, which was shown to regulate mono- and sesquiterpene biosynthesis in the type VI glandular trichomes of tomato leaves and stems (Spyropoulou et al., 2014; Xu et al., 2018), but that has not been implicated in the regulation of SGA biosynthesis yet.

In this study, we demonstrate that tomato MYC2, together with its homolog MYC1, is instrumental for the basal, but high SGA content in tomato organs. These two clade IIIe bHLH TFs ensure constitutive SGA biosynthesis by redundantly controlling basal expression of both cholesterol and SGA pathway genes. Although MYC-regulated specialized metabolism is typically induced following an elevation in intracellular JA-Ile levels, we found that the constitutive production of SGAs only partially relies on COI1-dependent JA signaling. Finally, CRISPR-Cas9(VQR) genome editing of an endogenous G-box, which is targeted by MYC1/2, in the promoter of the cholesterol biosynthesis gene *STEROL C-5(6) DESATURASE 2* (*C5-SD2*) leads to decreased constitutive expression of *C5-SD2*. This further supports the role of the tomato clade IIIe bHLH TFs in the regulation of constitutive SGA biosynthesis.

## RESULTS

### MYC2 Coordinates Constitutive Expression of SGA Biosynthesis Genes

To enhance our understanding of MYC2 as a regulator of SGA production, we consulted previously published RNA-seq data from wild-type and *MYC2-RNAi* seedlings (cultivar M82) (Du et al., 2017). Among the differentially expressed genes (DEGs) between mock-treated wild-type and *MYC2-RNAi* plants (false discovery rate [FDR]-adjusted P value < 0.05), we encountered genes known to be involved in the biosynthesis of SGAs and their precursors (Table 1). Indeed, transcript levels of genes encoding enzymes for cycloartenol, cholesterol, and SGA biosynthesis were all significantly reduced in mock-treated *MYC2-RNAi* seedlings compared with wild-type seedlings (Table 1), indicating that MYC2 helps ensure their basal expression.

**Table 1.** Genes Differentially Expressed Between Mock-Treated Wild-Type and *MYC2-RNAi* Tomato Seedlings That Belong to bHLH Clade IIIe or Are Responsible for SGA Biosynthesis

| SolycID | Gene Name | Log<sub>2</sub> Fold Change<sup>a</sup> | Adjusted P Value |
| --- | --- | --- | --- |
| bHLH Clade IIIe Transcription Factors: |  |  |  |
| Solyc08g076930.1.1 | *MYC2* | -1.33 | $1.28 \times 10^{-31}$ |
| Solyc08g005050.2.1 | *MYC1* | -0.45 | $3.82 \times 10^{-2}$ |
| Cholesterol and Phytosterol Biosynthesis Pathway: |  |  |  |
| Solyc04g070980.2.1 | *CAS* | -0.53 | $1.08 \times 10^{-9}$ |
| Solyc02g081730.2.1 | *3bHSD2* | -0.78 | $1.36 \times 10^{-7}$ |
| Solyc12g098640.1.1 | *CPI* | -0.44 | $3.75 \times 10^{-3}$ |
| Solyc01g008110.2.1 | *CYP51* | -0.34 | $6.91 \times 10^{-6}$ |
| Solyc09g009040.2.1 | *C14-R* | -0.55 | $4.35 \times 10^{-5}$ |
| Solyc06g082980.2.1 | *8,7-SI* | -0.98 | $2.50 \times 10^{-24}$ |
| Cholesterol Biosynthesis Pathway: |  |  |  |
| Solyc02g069490.2.1 | *SSR2* | -1.07 | $1.79 \times 10^{-8}$ |
| Solyc01g091320.2.1 | *SMO3* | -1.20 | $2.13 \times 10^{-19}$ |
| Solyc06g005750.2.1 | *SMO4* | -0.85 | $1.09 \times 10^{-15}$ |
| Solyc02g086180.2.1 | *C5-SD2* | -1.60 | $6.10 \times 10^{-66}$ |
| Solyc06g074090.2.1 | *7-DR2* | -0.97 | $3.09 \times 10^{-9}$ |
| SGA Biosynthesis Pathway: |  |  |  |
| Solyc07g043420.2.1 | *GAME11* | -1.39 | $3.93 \times 10^{-139}$ |
| Solyc07g043460.2.1 | *GAME6* | -1.52 | $5.27 \times 10^{-69}$ |
| Solyc12g006460.1.1 | *GAME4* | -0.70 | $4.63 \times 10^{-12}$ |
| Solyc12g006470.1.1 | *GAME12* | -1.15 | $2.25 \times 10^{-106}$ |
| Solyc01g073640.2.1 | *GAME25* | -0.87 | $3.67 \times 10^{-46}$ |
| Solyc07g043490.1.1 | *GAME1* | -1.13 | $8.10 \times 10^{-21}$ |
| Solyc07g043480.1.1 | *GAME17* | -0.89 | $5.49 \times 10^{-14}$ |
| Solyc07g043500.1.1 | *GAME18* | -1.31 | $8.40 \times 10^{-20}$ |
| RNA-seq data was previously published in (Du et al., 2017). |  |  |  |
| <sup>a</sup>Mean expression ratios calculated from three biologically independent experiments. |  |  |  |

To further investigate the potential role of MYC2 as a transcriptional activator of constitutive SGA biosynthesis, we generated three independent *myc2* loss-of-function hairy root lines (cultivar Moneymaker) using Clustered Regularly Interspaced Short Palindromic Repeats (CRISPR)-CRISPR associated protein 9 (CRISPR-Cas9) genome editing (Figure 1A and Supplemental Figure 1). We selected nine genes from the list of DEGs involved in SGA biosynthesis (Table 1) and measured their expression by quantitative real-time PCR (qPCR) in mock- and JA-treated control and *myc2* lines. Expression of several of these genes was significantly reduced in mock-treated *myc2* hairy root lines compared with control lines (Figure 1B) while their JA inducibility, however, remained intact (Supplemental Figure 2). Expression of *GAME9*, a known regulator of SGA biosynthesis, was comparable between mock-treated control and *myc2* root lines (Figure 1B). Targeted metabolite profiling showed there was only a small decrease in α-tomatine and dehydrotomatine levels between control and *myc2* hairy roots in both mock- and JA-treated conditions (Figure 1C). We did not observe a significant increase of these SGAs upon JA induction, in our hands, neither in control lines nor in *myc2* lines. Taken together, these observations suggest that besides MYC2, another JA-regulated TF is involved in the regulation of constitutive SGA production.

**Figure 1.**
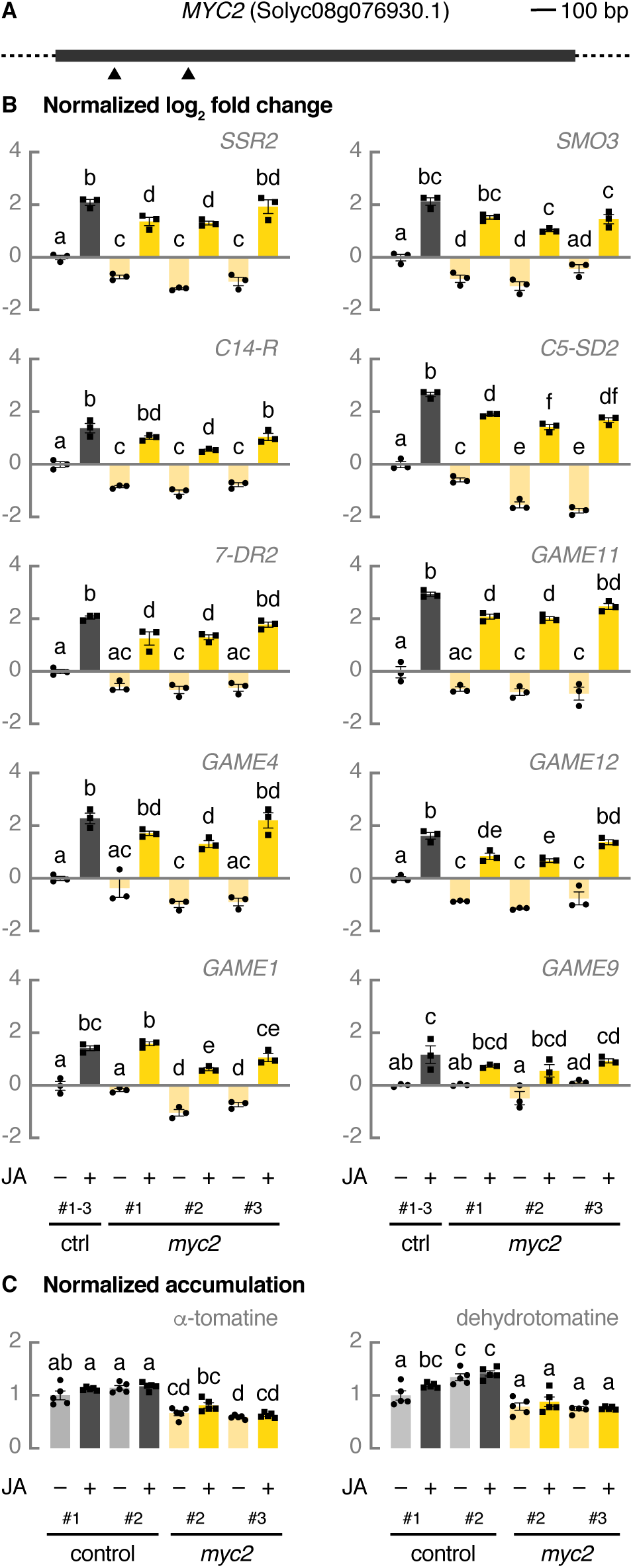
MYC2 coordinates constitutive SGA biosynthesis in tomato. (A) Schematic representation of *MYC2* with location of the CRISPR-Cas9 cleavage sites. The dark grey box represents the exon. Cas9 cleavage sites for two guide RNAs are indicated with arrowheads. (B) Relative expression of cholesterogenesis genes, SGA biosynthesis genes, and *GAME9* analyzed by qPCR. Control hairy root lines expressing *pCaMV35S::GUS* (grey bars) and *myc2* lines (yellow bars) were treated for 24 h with 50 μM of JA or an equal amount of ethanol. For control samples, cDNA of three biological replicates was pooled per independent line and treatment. Bars represent mean log_2_-transformed fold changes relative to the mean of three independent mock-treated control lines. Error bars denote standard error (n = 3). Individual mock- (●) and JA-treated (■) values are shown. Statistical significance was determined by ANOVA followed by Tukey post-hoc analysis (P<0.05; indicated by different letters). (C) Relative accumulation of *α*-tomatine and dehydrotomatine analyzed by LC-MS. Control hairy root lines expressing *pCaMV35S::GUS* (grey bars) and *myc2* lines (yellow bars) were treated for 24 h with 50 μM of JA or an equal amount of ethanol. Bars represent mean fold changes relative to the mean of mock-treated control^#1^. Error bars denote standard error (n=5). Individual mock- (●) and JA-treated (■) values are shown. Statistical significance was determined by ANOVA followed by Tukey post-hoc analysis (P<0.05; indicated by different letters). Abbreviations: *SSR2*, *STEROL SIDE CHAIN REDUCTASE 2*; *SMO3*, *C-4 STEROL METHYL OXIDASE 3*; *C14-R*, *STEROL C-14 REDUCTASE*; *7-DR2*, *7-DEHYDROCHOLESTEROL REDUCTASE 2*; *GUS*, *β-glucuronidase*.

For comparison, we measured the expression of a gene involved in JA signaling, *JAZ1* (Sun et al., 2011), and a gene involved in the biosynthesis of polyamines, *ORNITHINE DECARBOXYLASE* (*ODC*) (Acosta et al., 2005), which were previously reported to be upregulated upon JA treatment (Chen et al., 2006; Chini et al., 2017). In contrast to the downregulation of several SGA pathway genes in mock-treated *myc2* lines (Figure 1B), the expression of *JAZ1* and *ODC* was not reduced in mock-treated *myc2* lines compared to control lines (Figure 2A). The JA inducibility of *JAZ1* and *ODC* transcription in *myc2* hairy roots was not affected either (Figure 2A). Next, we determined the levels of two phenylpropanoid-polyamine conjugates, tris(dihydrocaffeoyl)spermine and *N*-caffeoylputrescine, which are both reported to be highly JA-inducible compounds (Chen et al., 2006). While the levels of two main SGAs were decreased between mock-treated control and *myc2* hairy roots (Figure 1C), the levels of tris(dihydrocaffeoyl)spermine and *N*-caffeoylputrescine were not (Figure 2B). JA-treated *myc2* lines, however, accumulated less of these polyamines than JA-treated control lines did (Figure 2B). These results suggest that MYC2 specifically regulates the constitutive biosynthesis of SGAs and not of other specialized metabolites such as the phenylpropanoid-polyamine conjugates. Conversely, MYC2 is clearly involved, together with other, yet elusive, TFs, in the induction of phenylpropanoid-polyamine conjugate biosynthesis upon JA signaling.

**Figure 2.**
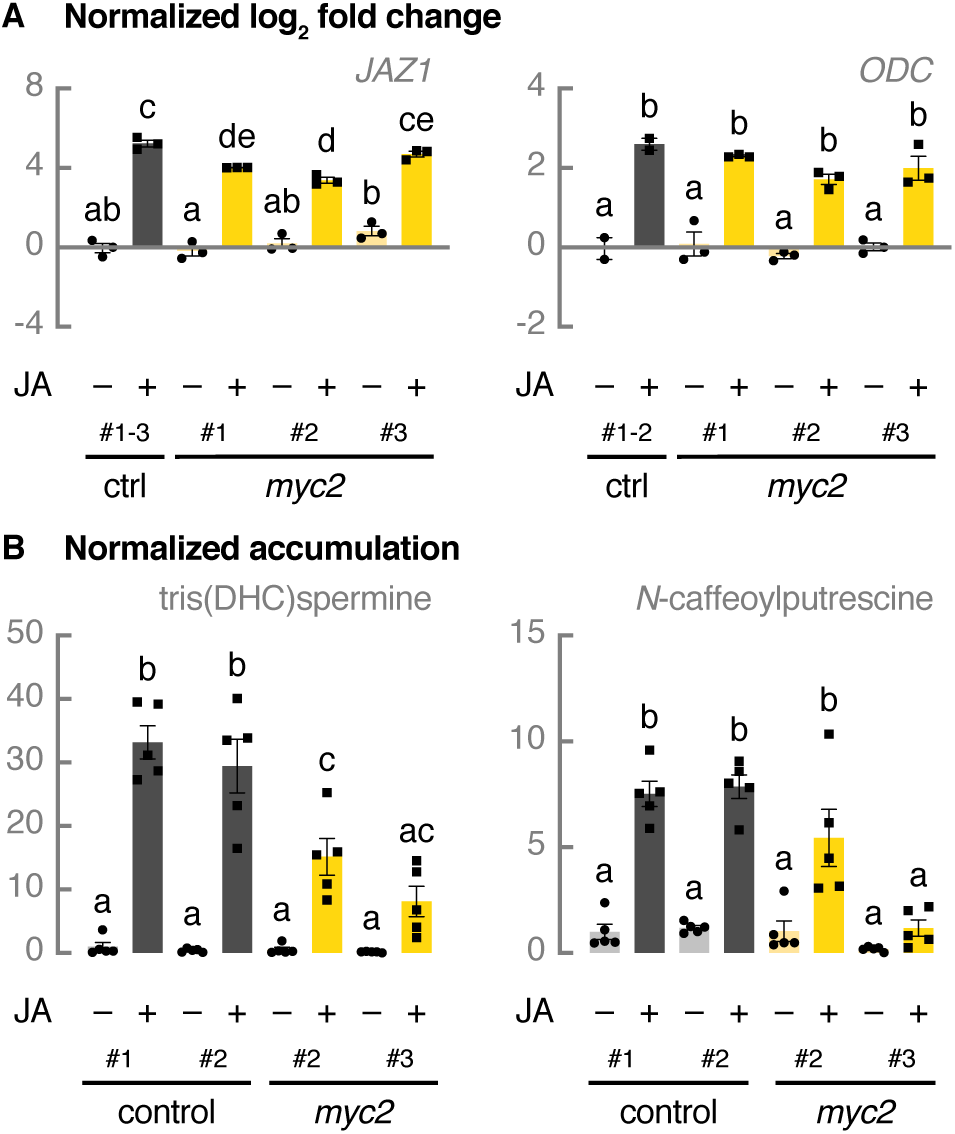
MYC2 helps ensure JA-induced polyamine production in tomato. (A) Relative expression of *JAZ1* and *ODC* analyzed by qPCR. Control hairy root lines expressing *pCaMV35S::GUS* (grey bars) and *myc2* lines (yellow bars) were treated for 24 h with 50 μM of JA or an equal amount of ethanol. For control samples, cDNA of three biological replicates was pooled per independent line and treatment. Bars represent mean log_2_-transformed fold changes relative to the mean of three independent mock-treated control lines. Error bars denote standard error (n = 3). Individual mock- (●) and JA-treated (■) values are shown. Statistical significance was determined by ANOVA followed by Tukey post-hoc analysis (P<0.05; indicated by different letters). (B) Relative accumulation of tris(dihydrocaffeoyl)spermine and *N*-caffeoylputrescine analyzed by LC-MS. Control hairy root lines expressing *pCaMV35S::GUS* (grey bars) and *myc2* lines (yellow bars) were treated for 24 h with 50 μM of JA or an equal amount of ethanol. Bars represent mean fold changes relative to the mean of mock-treated control^#1^. Error bars denote standard error (n=5). Individual mock- (●) and JA-treated (■) values are shown. Statistical significance was determined by ANOVA followed by Tukey post-hoc analysis (P<0.05; indicated by different letters). Abbreviations: DHC, dihydrocaffeoyl; *GUS*, *β-glucuronidase*.

### Tomato Has Two bHLH Clade IIIe Family Members

The intact JA inducibility but reduced basal expression of SGA pathway genes and the slight decrease of two main SGAs in *myc2* hairy root lines (Supplemental Figure 2 and Figure 1B–C) suggested that another JA-regulated TF is involved in the regulation of SGA biosynthesis. In Arabidopsis, bHLH clade IIIe consists of MYC2 and three other MYC TFs that all contribute to the JA response (Goossens et al., 2017). Therefore, using the PLAZA comparative genomics platform (Van Bel et al., 2018), we searched for other tomato orthologs of Arabidopsis MYC2 and only identified MYC1, a regulator of type VI glandular trichome development and volatile terpene biosynthesis within trichomes (Spyropoulou et al., 2014; Xu et al., 2018). Phylogenetic analysis revealed that duplication of an Arabidopsis *MYC2* ortholog must have occurred at the base of the Solanaceae and that both copies were retained in the *S. lycopersicum* genome (Figure 3A) as is the case with several other Solanaceae species (Xu et al., 2017). While the Arabidopsis bHLH clade IIIe consists of MYC2, MYC3, MYC4, and MYC5, it appears to be limited to only two members in Solanaceae species (Figure 3A). A survey of public transcriptome data (cultivar Heinz 1706) (Zouine et al., 2017) showed that the two tomato clade IIIe bHLH members, *MYC1* and *MYC2*, exhibit a similar expression pattern (Figure 3B). In addition, transcript levels of both of these TFs were increased upon JA treatment of control hairy roots (Figure 3C). Although MYC1 and MYC2 may have distinct roles, such as in the regulation of glandular trichome development and volatile terpene biosynthesis, which specifically involves MYC1 but not MYC2 (Spyropoulou et al., 2014; Xu et al., 2018), our data indicate that MYC1 and MYC2 might have overlapping functions as well.

**Figure 3.**
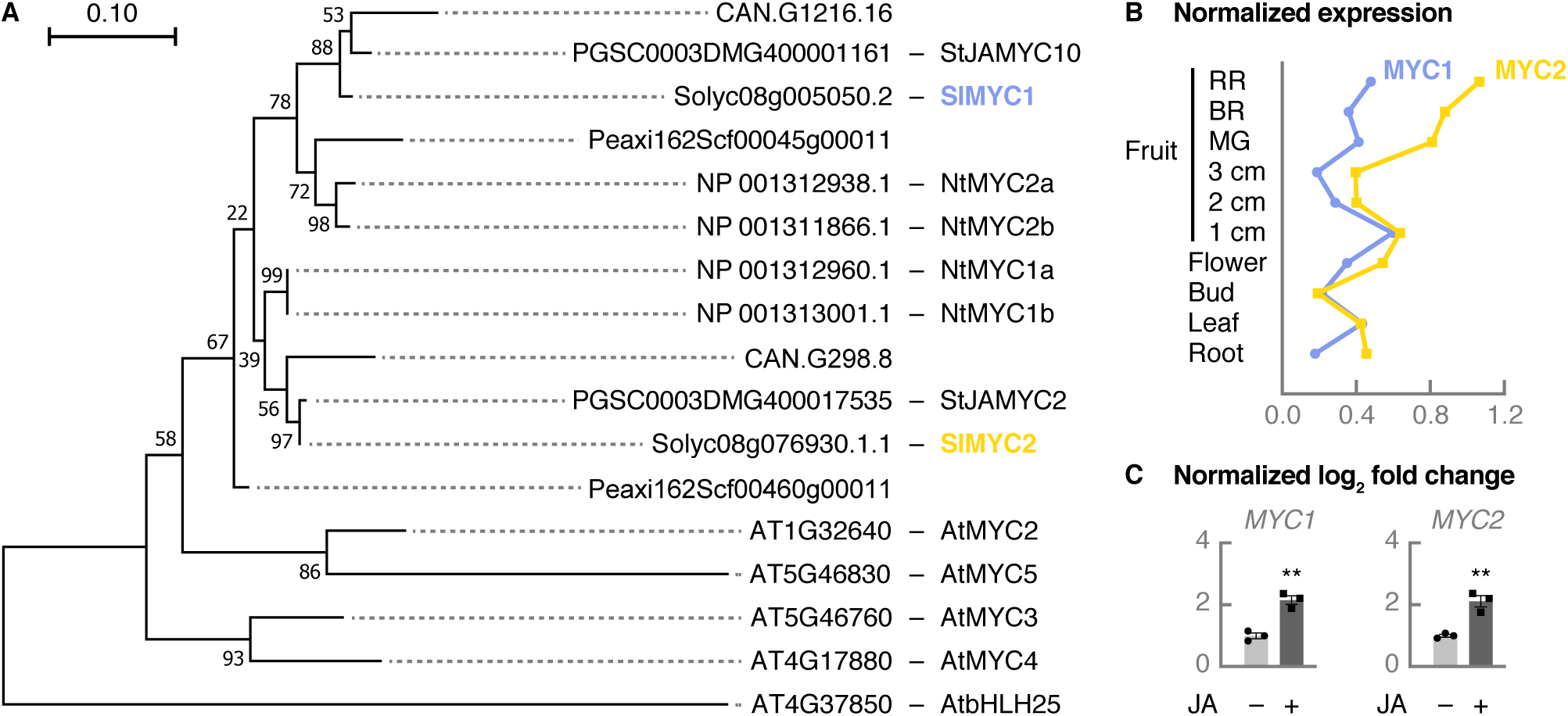
Tomato has two clade IIIe bHLH family members. (A) Phylogenetic analysis of *Arabidopsis thaliana* and Solanaceae bHLH clade IIIe members using MUSCLE and the Maximum Likelihood method. The tree is drawn to scale, with branch lengths measured in the number of substitutions per site. The analysis involved amino acid sequences from *Capsicum annuum* (CAN.G1216.16, CAN.G298.8), *Solanum tuberosum* (PGSC0003DMG400001161, PGSC0003DMG400017535), *S. lycopersicum* (Solyc08g005050.2, Solyc08g076930.1.1), *Petunia axillaris* (Peaxi162Scf00045g00011, Peaxi162Scf00460g00011), *Nicotiana tabacum* (NP_001312938.1, NP_001311866.1, NP_001312960.1, NP_001313001.1), and *A. thaliana* (AT1G32640, AT5G46830, AT5G46760, AT4G17880). The *A. thaliana* bHLH clade Iva member bHLH25 (AT4G37850) was used to root the tree. Numbers shown are bootstrap values in percentages (based on 1,000 replicates). Evolutionary analyses were conducted in MEGA7 (Kumar et al., 2016). (B) Normalized *MYC1* and *MYC2* expression profiles in different organs and developmental stages (cultivar Heinz 1706). Expression data were obtained from TomExpress (Zouine et al., 2017). (C) Relative expression of *MYC1* and *MYC2* analyzed by qPCR. Control hairy root lines expressing *pCaMV35S::GUS* were treated for 24 h with 50 μM of JA or an equal amount of ethanol. The cDNA of three biological replicates was pooled per independent line and treatment. Bars represent mean log_2_-transformed fold changes relative to the mean of three independent mock-treated control lines. Error bars denote standard error (n=3). Individual mock- (●) and JA-treated (■) values are shown. Statistical significance was determined by unpaired Student’s *t*-tests (**, P<0.01). Abbreviations: 1 cm, 1 cm immature green fruit; 2 cm, 2 cm immature green fruit; 3 cm, 3 cm immature green fruit; MG, mature green fruit; BR, breaker fruit; RR, red ripe (breaker + 10 days) fruit.

### MYC1 and MYC2 Redundantly Control SGA Biosynthesis

Previously, we reported that MYC2 and GAME9 act synergistically to transactivate the promoter of *C5-SD2*, a gene involved in cholesterogenesis, when fused to the *FIREFLY LUCIFERASE* (*fLUC*) gene in tobacco (*Nicotiana tabacum*) protoplasts (Cárdenas et al., 2016). Here, we show that a similar cooperative action can be observed between MYC1 and GAME9 (Figure 4A). A G-box present within the *C5-SD2* promoter was formerly found to be bound by MYC2 and to be essential for the transactivation of this promoter by the combined action of MYC2 and GAME9 (Cárdenas et al., 2016). Mutating that same G-box in a 333-bp *C5-SD2* promoter region, which was shown to be sufficient for synergistic transactivation by MYC1/MYC2 and GAME9 (Figure 4B), led to severely reduced induction of the luciferase activity (Figure 4C), while no luciferase induction was measured for a promoter deletion construct lacking the G-box (Figure 4C). Thus, these data indicate that MYC1 might play a similar role as MYC2 in regulating SGA biosynthesis by binding the G-box in the promoters of SGA pathway genes and activating their expression in synergy with GAME9.

**Figure 4.**
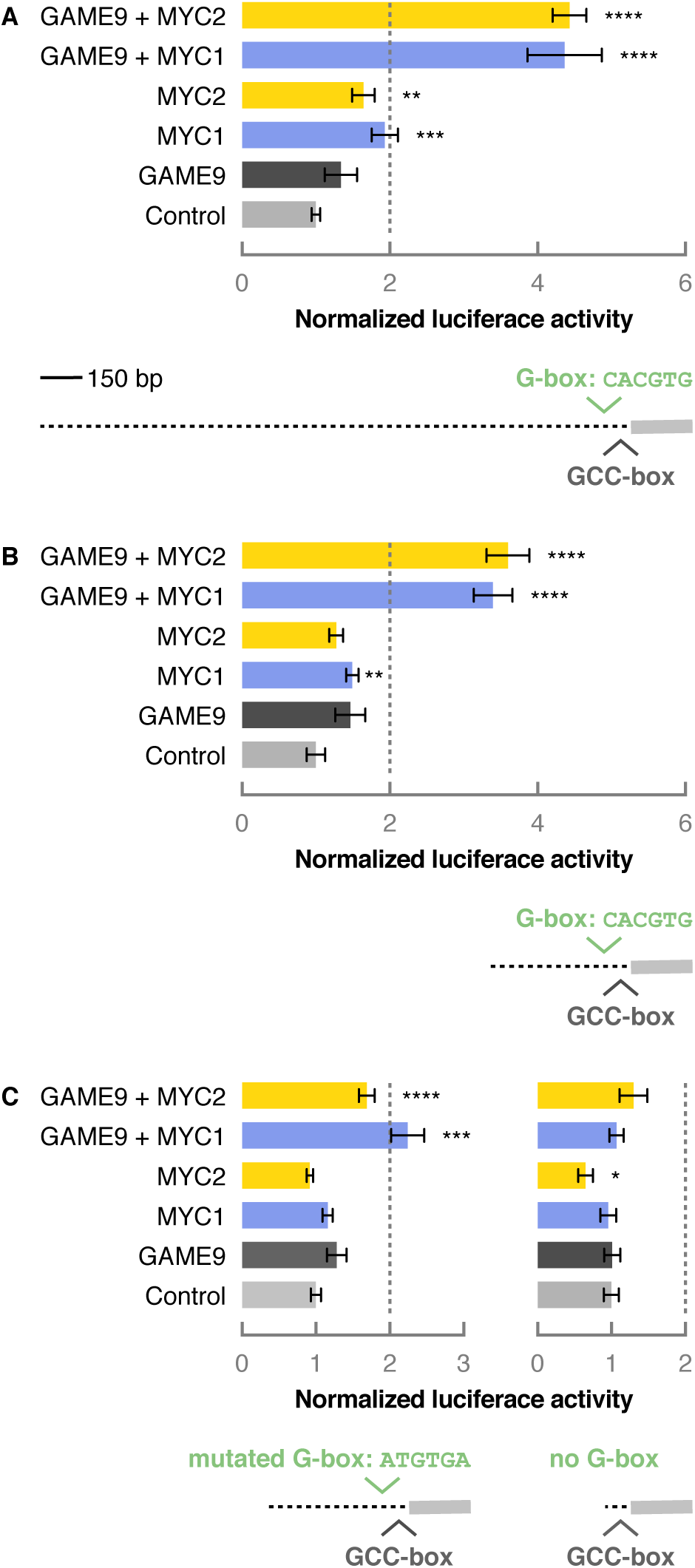
MYC1 and MYC2 transactivate the promoter of *C5-SD2* together with GAME9. (A-C) Tobacco BY-2 protoplasts were transfected with a *pC5-SD2(1549 bp)::fLUC* (A), *pC5-SD2(333 bp)::fLUC* (B), *pC5-SD2(333 bp with mutated G-box)::fLUC* or *pC5-SD2(207 bp without G-box)::fLUC* (C) reporter construct and effector constructs overexpressing *MYC1*, *MYC2*, *GAME9* or a combination thereof. A *pCaMV35S::rLUC* construct was co-transfected for normalization of fLUC activity. Bars represent mean fold changes relative to the mean of protoplasts transfected with a *pCaMV35S::GUS* control construct (grey bar). Dashed lines represent the 2-fold cut off for promoter transactivation. Error bars denote standard error (n = 8). Statistical significance was determined by unpaired Student’s *t*-tests (*, P<0.05; **, P<0.01; ***, P<0.001; ****, P<0.0001). Abbreviations: BY-2, Bright Yellow-2; *pC5-SD2*, promoter of *C5-SD2*; *GUS*, *β-glucuronidase*.

To explore whether MYC1 is indeed a regulator of SGA production *in planta*, we generated three independent *myc1* loss-of-function hairy root lines (cultivar Moneymaker) using CRISPR-Cas9 genome editing (Figure 5A and Supplemental Figure 3). A qPCR analysis of cycloartenol, cholesterol, and SGA biosynthesis genes revealed that some of them were significantly downregulated in mock-treated *myc1* hairy roots (Figure 5B), whereas no effect was observed on their JA inducibility (Supplemental Figure 4). We checked if the lack of JA-inducible effects in *myc1* and *myc2* single mutants could be explained by a genetic compensation response (Ma et al., 2019), but we did not observe an upregulation of *MYC2* in *myc1* lines or vice versa (Supplemental Figure 5). Levels of α-tomatine and dehydrotomatine were not affected in *myc1* lines compared with control lines, neither in mock- nor in JA-treated conditions (Figure 5C). The expression of *JAZ1* and *ODC* was unaltered in both mock- and JA-treated *myc1* hairy roots compared to control hairy roots (Figure 6A) and the levels of tris(dihydrocaffeoyl)spermine and *N*-caffeoylputrescine were only reduced between JA-treated control and *myc1* root lines (Figure 6B). These results are comparable to those observed for *myc2* lines (Figure 1–2), indicating that MYC1 and MYC2 might have redundant roles in regulating the constitutive production of SGAs and their precursors.

**Figure 5.**
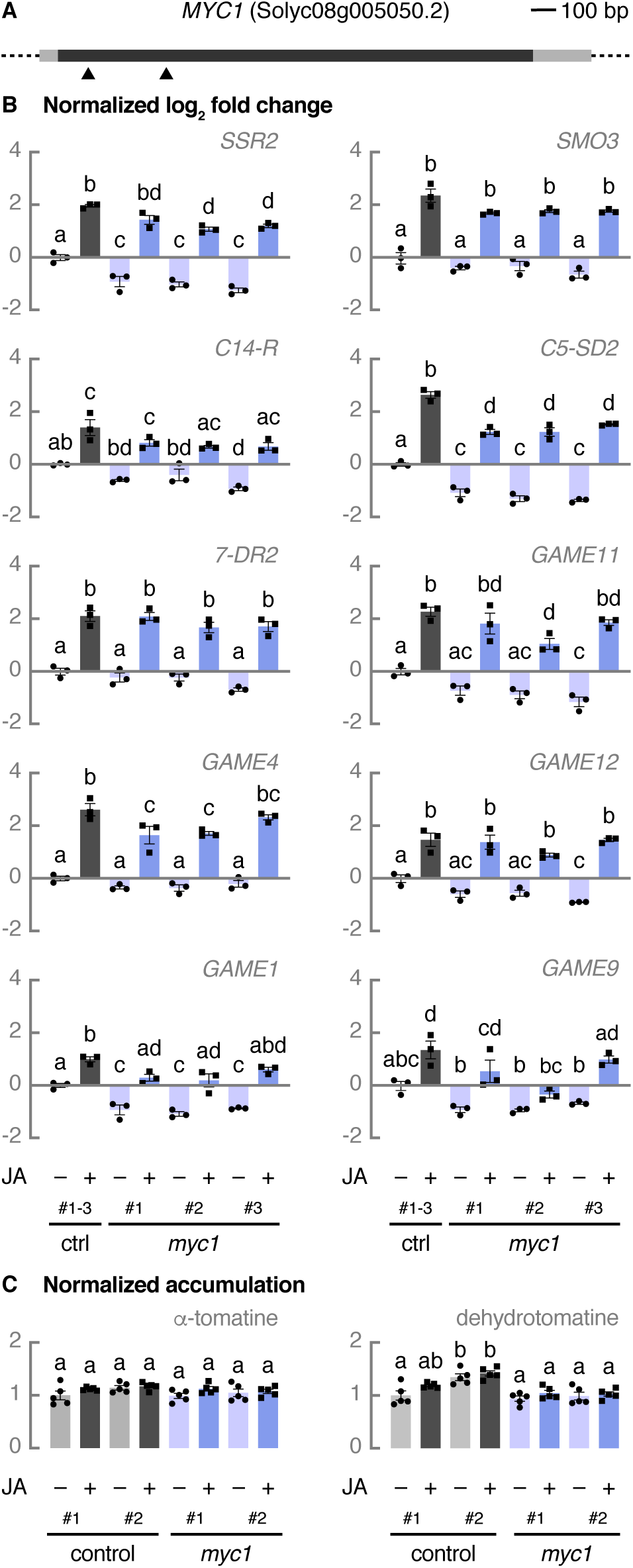
MYC1 regulates constitutive expression of SGA biosynthesis genes in tomato. (A) Schematic representation of *MYC1* with location of the CRISPR-Cas9 cleavage sites. The dark grey box represents the exon and light grey boxes represent the UTRs. Cas9 cleavage sites for two guide RNAs are indicated with arrowheads. (B) Relative expression of cholesterogenesis genes, SGA biosynthesis genes, and *GAME9* analyzed by qPCR. Control hairy root lines expressing *pCaMV35S::GUS* (grey bars) and *myc1* lines (blue bars) were treated for 24 h with 50 μM of JA or an equal amount of ethanol. For control samples, cDNA of three biological replicates was pooled per independent line and treatment. Bars represent mean log_2_-transformed fold changes relative to the mean of three independent mock-treated control lines. Error bars denote standard error (n = 3). Individual mock- (●) and JA-treated (■) values are shown. Statistical significance was determined by ANOVA followed by Tukey post-hoc analysis (P < 0.05; indicated by different letters). (C) Relative accumulation of *α*-tomatine and dehydrotomatine analyzed by LC-MS. Control hairy root lines expressing *pCaMV35S::GUS* (grey bars) and *myc1* lines (blue bars) were treated for 24 h with 50 μM of JA or an equal amount of ethanol. Bars represent mean fold changes relative to the mean of mock-treated control^#1^. Error bars denote standard error (n=5). Individual mock- (●) and JA-treated (■) values are shown. Statistical significance was determined by ANOVA followed by Tukey post-hoc analysis (P <0.05; indicated by different letters). Abbreviations: *SSR2*, *STEROL SIDE CHAIN REDUCTASE 2*; *SMO3*, *C-4 STEROL METHYL OXIDASE 3*; *C14-R*, *STEROL C-14 REDUCTASE*; *7-DR2*, *7-DEHYDROCHOLESTEROL REDUCTASE 2*; UTR, untranslated region; *GUS*, *β-glucuronidase*.

**Figure 6.**
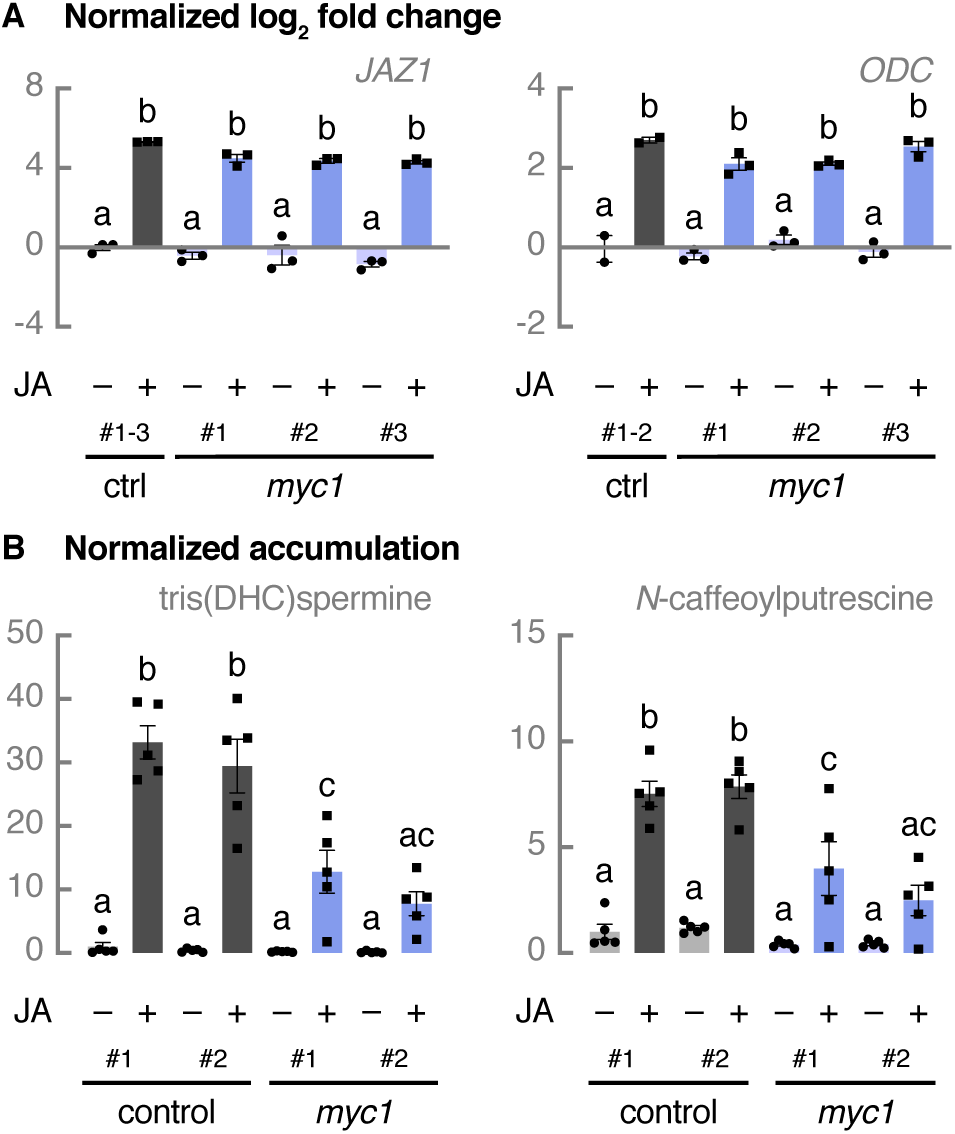
MYC1 helps ensure JA-induced polyamine biosynthesis in tomato. (A) Relative expression of *JAZ1* and *ODC* analyzed by qPCR. Control hairy root lines expressing *pCaMV35S::GUS* (grey bars) and *myc1* lines (blue bars) were treated for 24 h with 50 μM of JA or an equal amount of ethanol. For control samples, cDNA of three biological replicates was pooled per independent line and treatment. Bars represent mean log_2_-transformed fold changes relative to the mean of three independent mock-treated control lines. Error bars denote standard error (n = 3). Individual mock- (●) and JA-treated (■) values are shown. Statistical significance was determined by ANOVA followed by Tukey post-hoc analysis (P<0.05; indicated by different letters). (B) Relative accumulation of tris(dihydrocaffeoyl)spermine and *N*-caffeoylputrescine analyzed by LC-MS. Control hairy root lines expressing *pCaMV35S::GUS* (grey bars) and myc1 lines (blue bars) were treated for 24 h with 50 μM of JA or an equal amount of ethanol. Bars represent mean fold changes relative to the mean of mock-treated control^#1^. Error bars denote standard error (n=5). Individual mock- (●) and JA-treated (■) values are shown. Statistical significance was determined by ANOVA followed by Tukey post-hoc analysis (P<0.05; indicated by different letters). Abbreviations: DHC, dihydrocaffeoyl; *GUS*, *β-glucuronidase*.

Next, we used CRISPR-Cas9 genome editing to target *MYC1* and *MYC2* simultaneously, which yielded one double *myc1 myc2* hairy root knockout line (Supplemental Figure 6). We measured the expression of both SGA and more upstream biosynthesis genes and observed severely reduced transcript levels for most of them in mock-treated *myc1 myc2* hairy roots compared with control hairy roots (Figure 7A), while no decrease was detected in the expression of *JAZ1* and *ODC* (Figure 8A). Upon JA treatment, the transcription of not only cholesterol and SGA biosynthesis genes but also of *JAZ1* and *ODC* was not induced or not as strongly induced anymore in the *myc1 myc2* line compared with the control lines (Supplemental Figure 7 and Figure 8A). In accordance with these observations, the *myc1 myc2* line exhibited a 60–90% decrease in α-tomatine and dehydrotomatine content in both mock- and JA-treated conditions (Figure 7B). Although the biosynthesis of tris(dihydrocaffeoyl)spermine and *N*-caffeoylputrescine was no longer induced by JA in *myc1 myc2* hairy roots, their basal levels were not significantly reduced in *myc1 myc2* hairy roots compared to control hairy roots (Figure 8B). Thus, SGA pathway genes appear to be specifically affected while genes involved in canonical JA signaling and the biosynthesis of phenylpropanoid-polyamine conjugates are not. Taken together, these data demonstrate that MYC1 and MYC2 are functionally redundant in specifically controlling the constitutive biosynthesis of SGAs.

**Figure 7.**
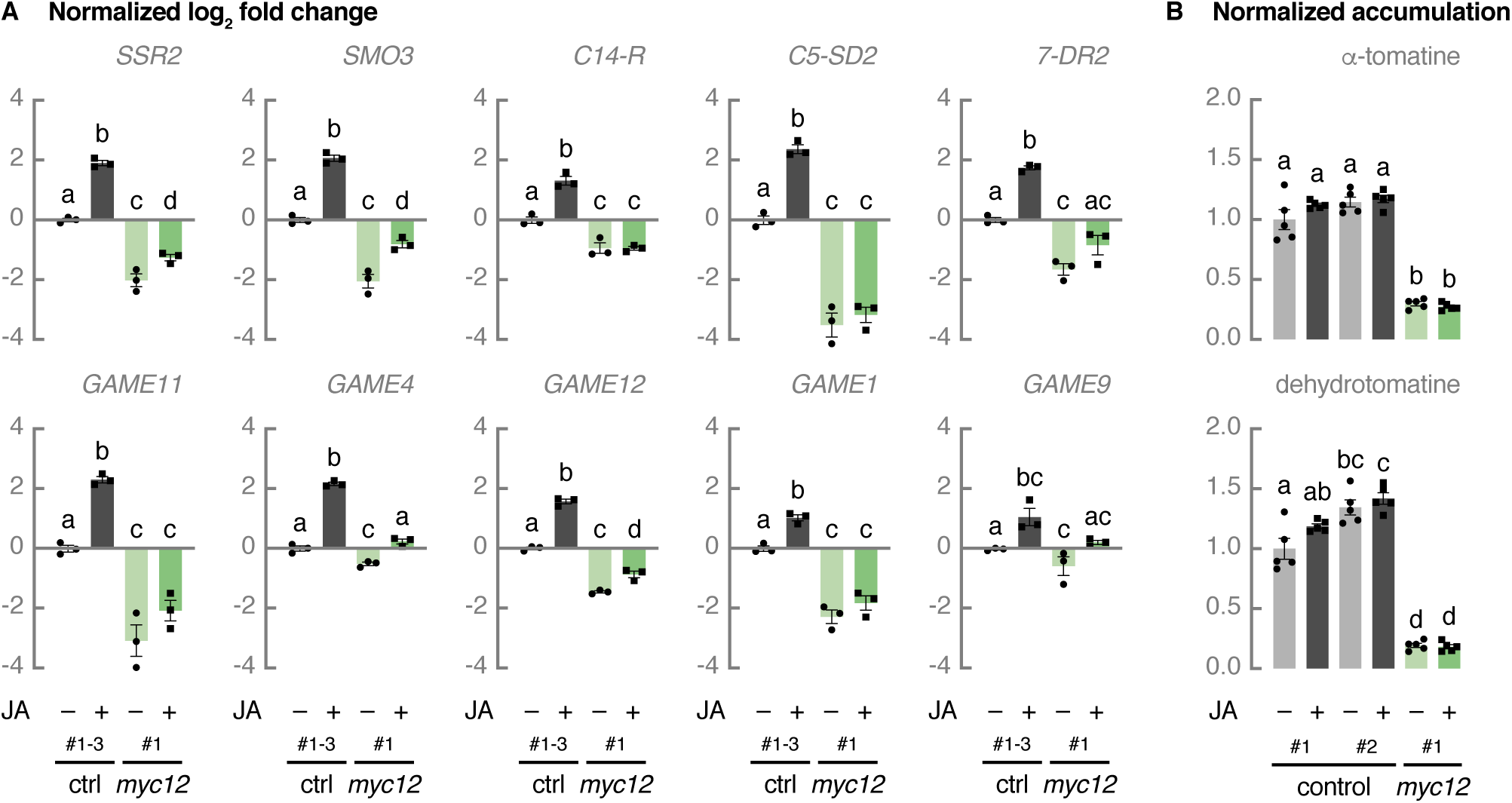
MYC1 and MYC2 redundantly regulate constitutive SGA biosynthesis in tomato. (A) Relative expression of cholesterogenesis genes, SGA biosynthesis genes, and *GAME9* analyzed by qPCR. Control hairy root lines expressing *pCaMV35S::GUS* (grey bars) and a *myc1 myc2* (*myc12*) line (green bars) were treated for 24 h with 50 μM of JA or an equal amount of ethanol. For control samples, cDNA of three biological replicates was pooled per independent line and treatment. Bars represent mean log_2_-transformed fold changes relative to the mean of three independent mock-treated control lines. Error bars denote standard error (n=3). Individual mock- (●) and JA-treated (■) values are shown. Statistical significance was determined by ANOVA followed by Tukey post-hoc analysis (P<0.05; indicated by different letters). (B) Relative accumulation of *α*-tomatine and dehydrotomatine analyzed by LC-MS. Control hairy root lines expressing *pCaMV35S::GUS* (grey bars) and a *myc1 myc2* (*myc12*) line (green bars) were treated for 24 h with 50 μM of JA or an equal amount of ethanol. Bars represent mean fold changes relative to the mean of mock-treated control^#1^. Error bars denote standard error (n = 5). Individual mock- (●) and JA-treated (■) values are shown. Statistical significance was determined by ANOVA followed by Tukey post-hoc analysis (P < 0.05; indicated by different letters). Abbreviations: *SSR2*, *STEROL SIDE CHAIN REDUCTASE 2*; *SMO3*, *C-4 STEROL METHYL OXIDASE 3*; *C14-R*, *STEROL C-14 REDUCTASE*; *7-DR2*, *7-DEHYDROCHOLESTEROL REDUCTASE 2*; *GUS*, *β-glucuronidase*.

**Figure 8.**
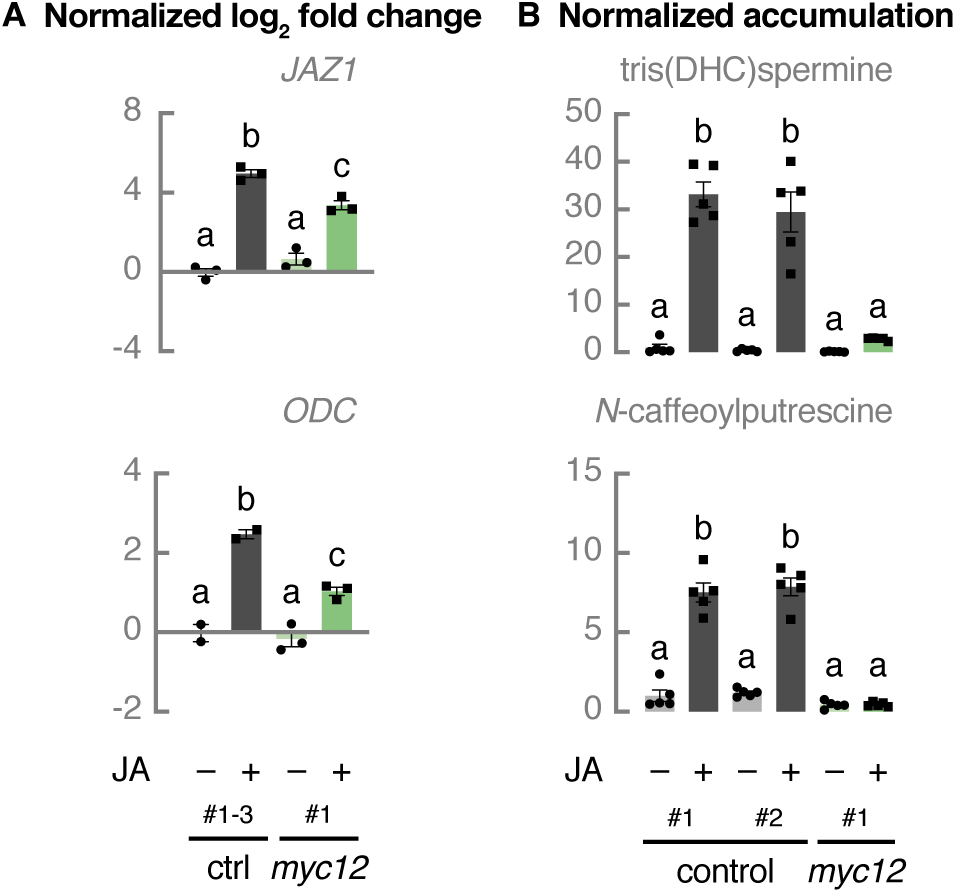
MYC1 and MYC2 redundantly regulate JA-induces polyamine accumulation in tomato. (A) Relative expression of *JAZ1* and *ODC* analyzed by qPCR. Control hairy root lines expressing *pCaMV35S::GUS* (grey bars) and a *myc1 myc2* (*myc12*) line (green bars) were treated for 24 h with 50 μM of JA or an equal amount of ethanol. For control samples, cDNA of three biological replicates was pooled per independent line and treatment. Bars represent mean log_2_-transformed fold changes relative to the mean of three independent mock-treated control lines. Error bars denote standard error (n = 3). Individual mock- (●) and JA-treated (■) values are shown. Statistical significance was determined by ANOVA followed by Tukey post-hoc analysis (P<0.05; indicated by different letters). (B) Relative accumulation of tris(dihydrocaffeoyl)spermine and *N*-caffeoylputrescine analyzed by LC-MS. Control hairy root lines expressing *pCaMV35S::GUS* (grey bars) and a *myc1 myc2* (*myc12*) line (green bars) were treated for 24 h with 50 μM of JA or an equal amount of ethanol. Bars represent mean fold changes relative to the mean of mock-treated control^#1^. Error bars denote standard error (n=5). Individual mock- (●) and JA-treated (■) values are shown. Statistical significance was determined by ANOVA followed by Tukey post-hoc analysis (P<0.05; indicated by different letters). Abbreviations: DHC, dihydrocaffeoyl; *GUS*, *β-glucuronidase*.

### SGA Biosynthesis Partially Depends on COI1-Mediated JA Signaling

The primary JA signaling pathway is highly conserved in the plant kingdom (Chini et al., 2016) and is initiated by the perception of JA-Ile by a co-receptor complex consisting of the F-box protein COI1 and a JAZ repressor (Yan et al., 2009; Sheard et al., 2010; Yan et al., 2018). Subsequent proteasomal degradation of the interacting JAZ protein (Chini et al., 2007; Thines et al., 2007) releases JA-regulated TFs from repression by JAZ (Chini et al., 2007; Thines et al., 2007; Chini et al., 2016). As in all investigated plant species, also in tomato the JAZ proteins can interact with MYC2 and, thereby, repress the transcription of MYC2-regulated genes (Du et al., 2017). Accordingly, COI1-dependent signaling has been suggested to be essential for JA-induced upregulation of SGA pathway genes (Abdelkareem et al., 2017). To investigate whether COI1 activity is also involved or required for the constitutive production of SGAs, we generated three independent *coi1* loss-of-function hairy root lines (cultivar Moneymaker) using CRISPR-Cas9 genome editing (Figure 9A and Supplemental Figure 8). Basal expression of not only several SGA pathway genes but also of *JAZ1* and *ODC* was significantly reduced in *coi1* hairy roots compared with control hairy roots (Figure 9B and Figure 10A). As expected, the expression of SGA biosynthesis genes, *JAZ1*, and *ODC* was not induced when *coi1* lines were treated with JA (Supplemental Figure 9 and Figure 10A). Furthermore, the α-tomatine and dehydrotomatine content in *coi1* hairy roots was decreased by 30–50% compared with control lines in both JA- and mock-treated conditions (Figure 9C), while tris(dihydrocaffeoyl)spermine and *N*-caffeoylputrescine levels were only reduced in JA-treated conditions (Figure 10B). This evidence suggests a partial dependence of constitutive SGA production on COI1-dependent signaling, possibly due to the stabilization and accumulation of JAZ proteins that block MYC1/2 activity.

**Figure 9.**
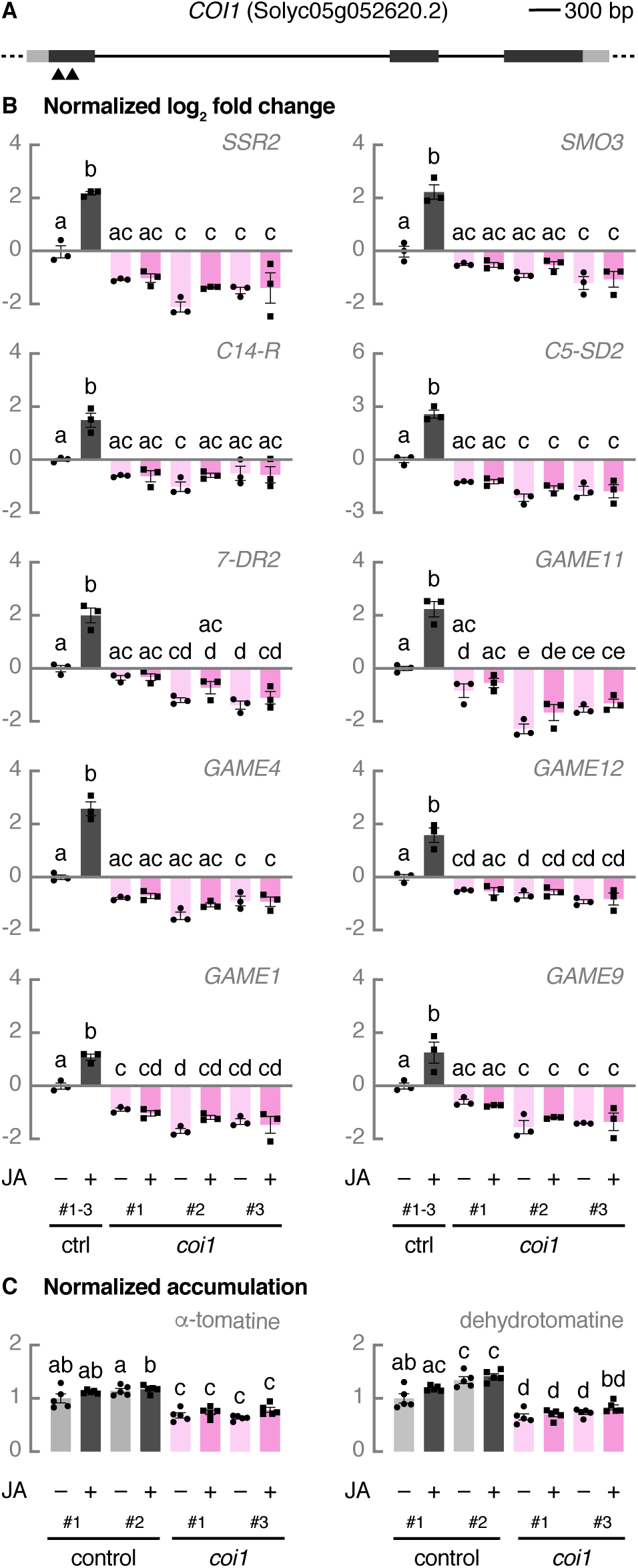
Constitutive SGA biosynthesis partially depends on COI1-mediated JA signaling. (A) Schematic representation of *COI1* with location of the CRISPR-Cas9 cleavage sites. Dark grey boxes represent exons, solid lines represent introns, and light grey boxes represent UTRs. Cas9 cleavage sites for two guide RNAs are indicated with arrowheads. (B) Relative expression of cholesterogenesis genes, SGA biosynthesis genes, and *GAME9* analyzed by qPCR. Control hairy root lines expressing *pCaMV35S::GUS* (grey bars) and *coi1* lines (pink bars) were treated for 24 h with 50 μM of JA or an equal amount of ethanol. For control samples, cDNA of three biological replicates was pooled per independent line and treatment. Bars represent mean log_2_-transformed fold changes relative to the mean of three independent mock-treated control lines. Error bars denote standard error (n=3). Individual mock- (●) and JA-treated (■) values are shown. Statistical significance was determined by ANOVA followed by Tukey post-hoc analysis (P < 0.05; indicated by different letters). (C) Relative accumulation of *α*-tomatine and dehydrotomatine analyzed by LC-MS. Control hairy root lines expressing *pCaMV35S::GUS* (grey bars) and *coi1* lines (pink bars) were treated for 24 h with 50 μM of JA or an equal amount of ethanol. Bars represent mean fold changes relative to the mean of mock-treated control^#1^. Error bars denote standard error (n=5). Individual mock- (●) and JA-treated (■) values are shown. Statistical significance was determined by ANOVA followed by Tukey post-hoc analysis (P<0.05; indicated by different letters). Abbreviations: *SSR2*, *STEROL SIDE CHAIN REDUCTASE 2*; *SMO3*, *C-4 STEROL METHYL OXIDASE 3*; *C14-R*, *STEROL C-14 REDUCTASE*; *7-DR2*, *7-DEHYDROCHOLESTEROL REDUCTASE 2*; UTR, untranslated region; *GUS*, *β-glucuronidase*.

**Figure 10.**
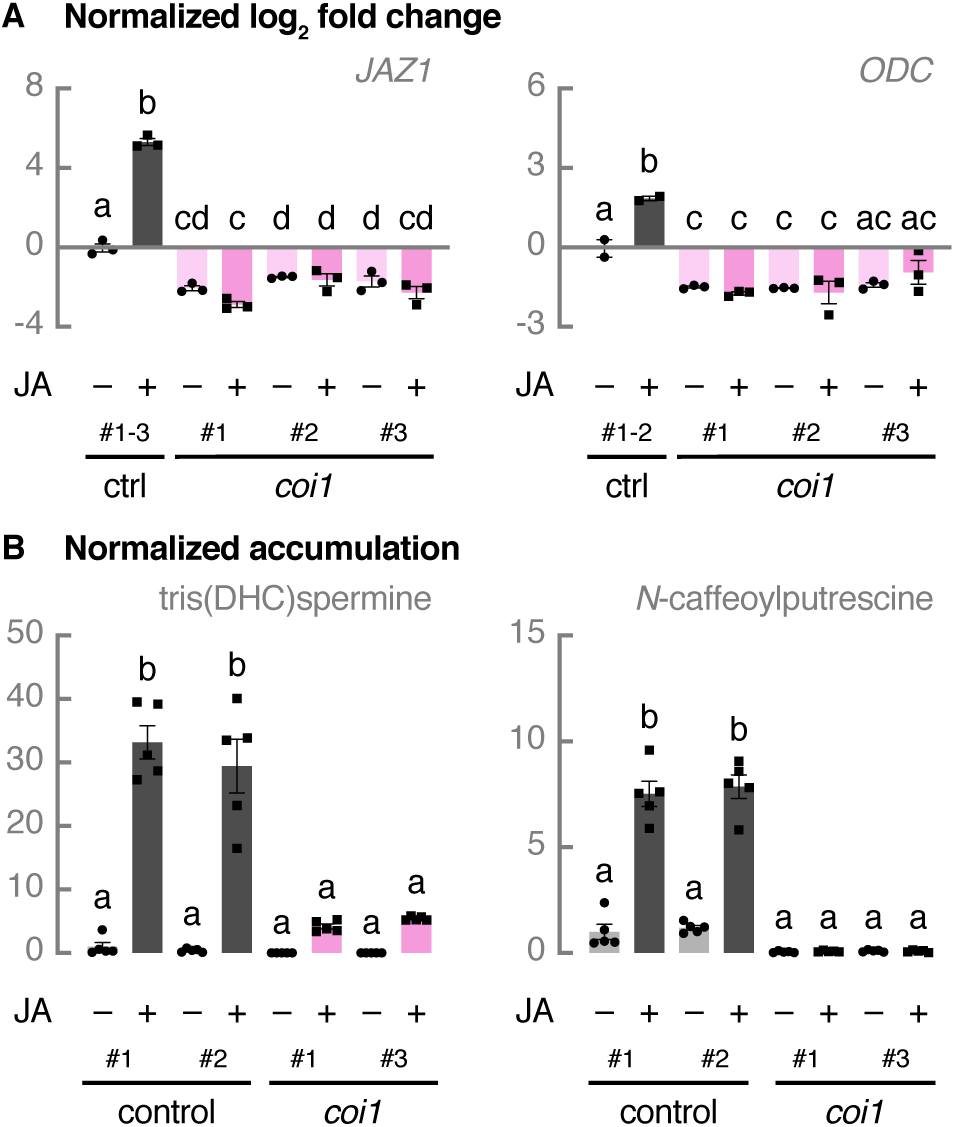
JA-induced polyamine biosynthesis depends on COI1-mediated JA signaling. (A) Relative expression of *JAZ1* and *ODC* analyzed by qPCR. Control hairy root lines expressing *pCaMV35S::GUS* (grey bars) and *coi1* lines (pink bars) were treated for 24 h with 50 μM of JA or an equal amount of ethanol. For control samples, cDNA of three biological replicates was pooled per independent line and treatment. Bars represent mean log_2_-transformed fold changes relative to the mean of three independent mock-treated control lines. Error bars denote standard error (n = 3). Individual mock- (●) and JA-treated (■) values are shown. Statistical significance was determined by ANOVA followed by Tukey post-hoc analysis (P<0.05; indicated by different letters). (B) Relative accumulation of tris(dihydrocaffeoyl)spermine and *N*-caffeoylputrescine analyzed by LC-MS. Control hairy root lines expressing *pCaMV35S::GUS* (grey bars) and *coi1* lines (pink bars) were treated for 24 h with 50 μM of JA or an equal amount of ethanol. Bars represent mean fold changes relative to the mean of mock-treated control^#1^. Error bars denote standard error (n=5). Individual mock- (●) and JA-treated (■) values are shown. Statistical significance was determined by ANOVA followed by Tukey post-hoc analysis (P<0.05; indicated by different letters). Abbreviations: DHC, dihydrocaffeoyl; *GUS*, *β-glucuronidase*.

### Genome Editing of a G-Box Decreases Constitutive *C5-SD2* Expression

In all investigated mutant genotypes (*myc1*, *myc2*, *myc1 myc2*, and *coi1*), the cholesterogenesis gene *C5-SD2* showed the strongest decrease in gene expression (Figure 1B, 5B, 7A, and 9B). Transient expression assays in tobacco protoplasts showed that a G-box present within the *C5-SD2* promoter is necessary for the transactivation of this promoter by MYC1 or MYC2 in combination with GAME9 (Figure 4) (Cárdenas et al., 2016). To investigate the relevance of this G-box *in planta,* we decided to target this *cis*-regulatory element in tomato hairy roots by genome editing using an engineered version of Cas9 that recognizes 5’-NGA-3’ as protospacer adjacent motif (PAM) (Kleinstiver et al., 2015). Three independent hairy root lines, denominated *g* lines, were generated in which the G-box motif was either deleted or disrupted. Line *g^#1^* contains a 33-bp deletion removing the G-box, while *g^#2^* and *g^#3^* carry a thymidine insertion and a 2-bp deletion within the G-box, respectively (Figure 11A). The 2-bp deletion in line *g^#3^* creates an alternative G-box (5’-CACGTT-3’). The transcript level of *C5-SD2* was significantly reduced in both mock- and JA-treated *g* lines compared with control lines, with the exception of mock-treated *g^#3^* hairy roots (Figure 11B). However, the JA-induced upregulation of *C5-SD2* in all three *g* lines was comparable to that in control hairy roots (Supplemental Figure 10), suggesting that the observed decrease in *C5-SD2* transcript levels in JA-treated conditions is merely due to the reduced basal *C5-SD2* expression. Next, we cloned *C5-SD2* promoter fragments from the *g* lines and fused them to the *fLUC* gene to create reporter constructs corresponding to the genome edited *C5-SD2* promoters. Transient expression assays in tobacco protoplasts showed that MYC2 and GAME9 were unable to transactivate these genome edited *C5-SD2* promoter fragments in which the G-box was either removed or disrupted (Figure 11C).

**Figure 11.**
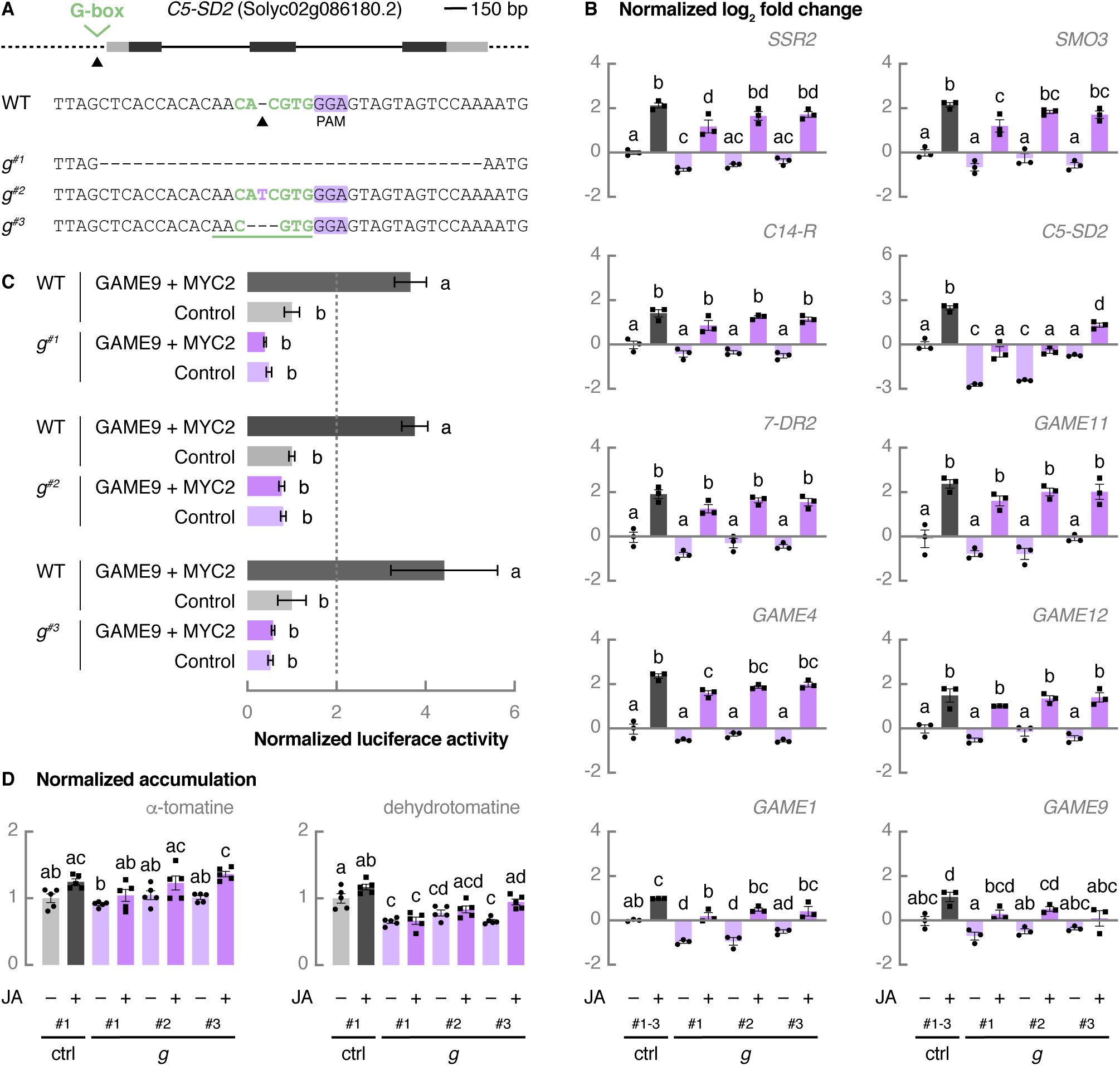
Genome editing of a G-box decreases constitutive *C5-SD2* expression. (A) Schematic representation of *C5-SD2* with location of the CRISPR-Cas9 cleavage site and G-box mutant sequences. Dark grey boxes represent exons, solid lines represent introns, and light grey boxes represent UTRs. The Cas9 cleavage site for the guide RNA targeting the G-box is indicated with an arrowhead. Sequences of three independent G-box mutant (*g*) lines are shown. G-box sequence is indicated in green font, the Cas9(VQR) PAM is marked in purple, inserted bases are shown in purple, deleted bases are replaced by a dash, and sequence gap length is shown between parentheses. Bases that make up an alternative G-box are green underlined. (B) Relative expression of cholesterogenesis genes, SGA biosynthesis genes, and *GAME9* analyzed by qPCR. Control hairy root lines expressing *pCaMV35S::GUS* (grey bars) and *g* lines (purple bars) were treated for 24 h with 50 μM of JA or an equal amount of ethanol. For control samples, cDNA of three biological replicates was pooled per independent line and treatment. Bars represent mean log_2_-transformed fold changes relative to the mean of three independent mock-treated control lines. Error bars denote standard error (n=3). Individual mock- (●) and JA-treated (■) values are shown. Statistical significance was determined by ANOVA followed by Tukey post-hoc analysis (P<0.05; indicated by different letters). (C) Tobacco BY-2 protoplasts were transfected with a *pC5-SD2(g^#1^; 1406 bp)::fLUC*, *pC5-SD2(g^#2^; 1406 bp)::fLUC* or *pC5-SD2(g^#3^; 1406 bp)::fLUC* reporter construct and effector constructs overexpressing *MYC2* and *GAME9*. A *pCaMV35S::rLUC* construct was co-transfected for normalization of fLUC activity. Bars represent mean fold changes relative to the mean of protoplasts transfected with a *pC5-SD2(WT; 1406 bp)::fLUC* reporter construct and a *pCaMV35S::GUS* control construct. Dashed lines represent the 2-fold cut off for promoter transactivation. Error bars denote standard error (n = 8). Statistical significance was determined by ANOVA followed by Tukey post-hoc analysis (P<0.05; indicated by different letters). (D) Relative accumulation of *α*-tomatine and dehydrotomatine analyzed by LC-MS. Control hairy root lines expressing *pCaMV35S::GUS* (grey bars) and *g* lines (purple bars) were treated for 24 h with 50 μM of JA or an equal amount of ethanol. Bars represent mean fold changes relative to the mean of mock-treated control^#1^. Error bars denote standard error (n=5). Individual mock- (●) and JA-treated (■) values are shown. Statistical significance was determined by ANOVA followed by Tukey post-hoc analysis (P<0.05; indicated by different letters). Abbreviations: *SSR2*, *STEROL SIDE CHAIN REDUCTASE 2*; *SMO3*, *C-4 STEROL METHYL OXIDASE 3*; *C14-R*, *STEROL C-14 REDUCTASE*; *7-DR2*, *7-DEHYDROCHOLESTEROL REDUCTASE 2*; UTR, untranslated region; PAM, protospacer adjacent motif; *GUS*, *β-glucuronidase*; BY-2, Bright Yellow-2; *pC5-SD2*, promoter of *C5-SD2*.

The expression levels of other cholesterol and most SGA biosynthesis genes were not altered in mock-treated *g* lines (Figure 11B), indicating that the effect on *C5-SD2* expression was specific and had no general feedback effects on other cholesterol and SGA pathway genes. Nonetheless, we observed a small but significant decrease in dehydrotomatine levels compared with control lines (Figure 11D), indicating that targeting of specific and essential *cis*-regulatory elements in promoters of key pathway genes is sufficient to alter metabolic fluxes in pathways and, thereby, modulate metabolite levels. For comparison, we measured the α-tomatine and dehydrotomatine content in two independent *c5-sd2* loss-of-function hairy root lines (cultivar Moneymaker) obtained by CRISPR-Cas9 genome editing (Supplemental Figure 11A). The levels of the main SGAs were reduced by approximately 50% in both mock- and JA-treated conditions compared with control lines (Supplemental Figure 11B). These results support our hypothesis that MYC1 and MYC2 help ensure constitutive *C5-SD2* expression by binding a G-box in the *C5-SD2* promoter.

## DISCUSSION

Throughout the plant kingdom, transcriptional regulators belonging to the clade IIIe bHLH TFs are master regulators of the JA-induced production of specialized metabolites that help plants fend off biotic enemies (De Geyter et al., 2012; Goossens et al., 2017). Here, we show that two tomato clade IIIe bHLH TFs, MYC1 and MYC2, control the constitutive production of SGAs that grant protection against a wide variety of herbivores and pathogens by making up a chemical defense barrier (Friedman, 2002). Accordingly, CRISPR-Cas9-mediated disruption or deletion of an endogenous G-box, which is targeted by MYC1/2, in the promoter of *C5-SD2* leads to decreased basal *C5-SD2* expression. Although the activation of specialized metabolism by MYC TFs is typically initiated by the perception of JA-Ile, constitutive SGA biosynthesis seems to only partially rely upon COI1-dependent signaling.

### JA-Regulated TFs Control Constitutive Alkaloid Production in Solanaceous Species

SGAs provide multiple members of the *Solanum* genus constitutive protection against a broad range of herbivores and pathogens (Friedman, 2002). Here, we report that the tomato JA-regulated TFs MYC1 and MYC2 coordinate the basal biosynthesis of these cholesterol-derived products. A double *myc1 myc2* hairy root knockout line displayed suppressed expression of genes known to be involved in the biosynthesis of SGAs and their precursors. In addition, the expression of several SGA pathway genes in *myc1 myc2* root lines was not induced anymore by JA treatment. Although *myc1 myc2* hairy roots no longer exhibited JA-induced upregulation of *ODC*, which encodes an enzyme in the highly JA-inducible polyamine pathway (Chen et al., 2006), the basal expression of *ODC* was unaffected. Accordingly, targeted metabolite profiling showed that *myc1 myc2* hairy roots contained severely reduced constitutive levels of the main tomato SGAs α-tomatine and dehydrotomatine but not of the phenylpropanoid-polyamine conjugates tris(dihydrocaffeoyl)spermine and *N*-caffeoylputrescine. The levels of the latter compounds in *myc1 myc2* root lines were only affected in JA-treated conditions. Basal transcription of some, but not all, cholesterol and SGA biosynthesis genes was reduced in single *myc1* and *myc2* knockout lines, however, their JA inducibility was retained. Moreover, only a modest decrease in α-tomatine and dehydrotomatine content was observed in *myc2* hairy root lines alone. This indicates that there is functional redundancy between MYC1 and MYC2 in the control of SGA accumulation. Likewise, both constitutive and insect-inducible glucosinolate production in Arabidopsis is redundantly controlled by the clade IIIe bHLH TFs MYC2, MYC3, and MYC4 (Schweizer et al., 2013).

Both MYC1 and MYC2 directly regulate the expression of *C5-SD2*, a cholesterogenesis gene, and likely of other cholesterol and SGA biosynthesis genes as well, in synergy with GAME9 by binding G-box and GC-rich elements in their promoters. Disruption or deletion of a G-box in the endogenous promoter of *C5-SD2* by genome editing leads to reduced *C5-SD2* transcription in both mock- and JA-treated conditions, which suggests that the synergistic action of these transcriptional regulators contributes to the basal expression of *C5-SD2*. Like our *myc1 myc2* hairy root knockout line, tomato plant lines in which *GAME9* is either silenced or mutated display suppressed basal transcript levels of cholesterol and SGA biosynthesis genes (Cárdenas et al., 2016; Nakayasu et al., 2018). Consequently, these lines accumulate less SGAs (Cárdenas et al., 2016; Nakayasu et al., 2018), suggesting that GAME9, another JA-regulated TF, regulates the basal expression of genes needed for the constitutive production of SGAs. Interestingly, tobacco MYC1 and MYC2 orthologs control the production of nicotine, another constitutively highly accumulating alkaloid, together with ERF189, a tobacco JA-regulated AP2-ERF family member related to GAME9 (Shoji and Hashimoto, 2011). Target gene promoters harbor G-box and GC-rich elements that allow binding of these clade IIIe bHLH TFs and ERF189, respectively (Shoji and Hashimoto, 2011; Kajikawa et al., 2017; Xu et al., 2017). Silencing of *MYC1* or *MYC2* orthologs in tobacco leads to suppressed constitutive transcription of genes involved in nicotine biosynthesis and severely reduced basal alkaloid levels (Shoji and Hashimoto, 2011). Furthermore, like GAME9, ERF189 controls alkaloid accumulation in unelicited conditions since *nic1 nic2* hairy roots, in which the most severely repressed AP2/ERF is *ERF189*, display decreased basal expression of nicotine biosynthesis and transport genes as well as declined constitutive alkaloid production (Shoji et al., 2010). This suggests that the role of clade IIIe bHLH TFs in the regulation of constitutive biosynthesis of bioactive specialized metabolites may occur within additional *Solanaceae* species and might even be widespread within the plant kingdom.

An important question that remains is how these JA-regulated TFs are able to drive constitutive biosynthesis of highly accumulating alkaloids in *Solanaceae* members. The proximity of G-box and GC-rich elements in their target promoters and the collaborative action of these clade IIIe bHLH and AP2/ERF TFs suggest their cooperative binding, which can be a way to enhance their specificity and binding affinity for *cis*-regulatory elements (CREs) (Brkljacic and Grotewold, 2017). Target specificity of Arabidopsis MYC2/MYC3/MYC4 has been proposed to be governed by their interaction with R2R3-MYB TFs (Schweizer et al., 2013). Furthermore, competitive binding between these MYBs and the JAZ repressors to the JAZ interaction domain of MYC2/3/4 has already been forwarded as a mechanism for the regulation of constitutive glucosinolate production in Arabidopsis (Schweizer et al., 2013). Thus, it is possible that tomato MYC1/2 and GAME9, as well as their tobacco counterparts, form protein complexes that may facilitate the shielding of clade IIIe bHLH TFs from JAZ repressors.

### Partial Dependence of Constitutive SGA Biosynthesis on JA Signaling

JA signaling provokes transcriptional reprograming, leading toward the biosynthesis of species-specific defense compounds across the plant kingdom. Although perception of JA-Ile typically promotes fast and strong upregulation of specialized metabolism, in our hands tomato hairy roots that were treated with JA for one day did not display a marked increase in SGA levels whereas they did in the levels of phenylpropanoid-polyamine conjugates. Only after three to four days of continuous JA treatment, we and others were able to observe a modest increase in SGA content of 1.6- to 1.8-fold (Supplemental Figure 12) (Nakayasu et al., 2018), suggesting that this may not be a primary effect of JA signaling. The same holds true for the limited JA-induced upregulation of nicotine biosynthesis in tobacco plants and hairy roots (Shoji et al., 2008). In both tomato and tobacco, COI1-mediated perception of JA-Ile has been suggested to be essential for the minimal increase in alkaloid production upon JA elicitation (Shoji et al., 2008; Abdelkareem et al., 2017). Here, we report that constitutive SGA biosynthesis declines in mock-treated tomato *coi1* loss-of-function mutants, which confirms previous observations (Abdelkareem et al., 2017). This might be due to the stabilization and accumulation of JAZ proteins that block the activity of MYC1/2. The decrease in constitutive SGA content in *coi1* lines, however, is not as severe as in the double *myc1 myc2* knockout line. Hence, this suggests that the regulation of basal SGA production only partially relies upon COI1-dependent JA signaling and that MYC1 and MYC2, likely together with GAME9, are able to regulate SGA biosynthesis independent of JA signaling as well. The reduction, but not absence, of SGAs in *spr2* tomato plants, in which JA biosynthesis is impaired, further supports this notion (Montero-Vargas et al., 2018).

Transcriptional coordination of genes involved in the same specialized metabolic pathway can be accomplished by their promoters acquiring CREs that can be bound by JA-regulated TFs (Mertens et al., 2016; Shoji, 2019). It is therefore plausible to assume that, through the recruitment of G-box and GCC-box elements to the promoters of alkaloid biosynthesis genes, the JA-regulated MYC1/2 and GAME9, and their orthologs in tobacco, evolved to accommodate the constitutive chemical defense barrier made up of alkaloids.

## METHODS

### DNA Constructs

#### Transient Expression Assay Constructs

For transient expression assays, the coding sequence of tomato *MYC1* was PCR-amplified with the primers listed in Supplemental Table 1 and recombined in a Gateway donor vector (Invitrogen). Subsequently, a Gateway LR reaction (Invitrogen) was performed with the p2GW7 vector (Vanden Bossche et al., 2013). The *C5-SD2* promoter regions in which a G-box was disrupted or removed were PCR-amplified from *g* hairy root lines (cultivar Moneymaker) and recombined in a Gateway donor vector (Invitrogen). Next, Gateway LR reactions (Invitrogen) were performed with the pGWL7 vector (Vanden Bossche et al., 2013). All other constructs used for transient expression assays were generated previously (Cárdenas et al., 2016).

#### CRISPR-Cas9 Constructs

To select CRISPR-Cas9 guide (g)RNA target sites, CRISPOR (http://crispor.tefor.net/) (Haeussler et al., 2016) was used, with as PAM requirement 5’-NGA-3’ for targeting the G-box in the *C5-SD2* promoter and 5’-NGG-3’ for single and double knockouts. CRISPR-Cas9 constructs were cloned as previously described (Fauser et al., 2014; Ritter et al., 2017; Pauwels et al., 2018). Briefly, for each gRNA target site, two complementary oligonucleotides with 4-bp overhangs (Supplemental Table 1) were annealed and inserted by a Golden Gate reaction with *Bpi*I (Thermo Scientific) and T4 DNA ligase (Thermo Scientific) in following Gateway entry vectors: pEN-C1.1 (Fauser et al., 2014) was used for targeting the G-box in the *C5-SD2* promoter by a single gRNA approach, pMR217 (L1–R5) and pMR218 (L5–L2) (Ritter et al., 2017) were used for single *myc1*, *myc2*, *coi1*, and *c5-sd2* knockouts by a dual gRNA approach, and pMR217 (L1–R5) (Ritter et al., 2017), pMR219 (L5–L4), pMR204 (R4–R3), and pMR205 (L5–L2) were used for the double *myc1 myc2* knockout. To allow combining four gRNA modules, primers (Supplemental Table 1) were designed to amplify the gRNA module from pEn-C1.1 (L1–L2) (Fauser et al., 2014) adding appropriate attB/attBr flanking sites (L5–L4, B4r–B3r, and L3–L2) to each fragment. The amplified fragments were then cloned into the corresponding pDONR221 vector (pDONR221 P5–P4, P4r–P3r, and P3–P2) by Gateway BP reactions (Invitrogen) to generate entry clones suitable for MultiSite Gateway LR cloning. An additional *Bbs*I site in the pDONR backbone was eliminated by site-directed mutagenesis using primers noBbsI_F and noBbsI_R (Supplemental Table 1) followed by an In-Fusion reaction (Takara Bio USA). In order to yield the final binary vectors, (MultiSite) Gateway LR reactions (Invitrogen) were used. One gRNA module was recombined with pDe-Cas9(VQR)-Km (Kleinstiver et al., 2015; Swinnen et al., 2020) to target the G-box in the *C5-SD2* promoter. Two and four gRNA modules were recombined with pDe-Cas9-Km (Ritter et al., 2017) for single *myc1*, *myc2*, *coi1*, and *c5-sd2* knockouts and a double *myc1 myc2* knockout, respectively.

### Transient Expression Assays in Tobacco Protoplasts

Transient expression assays in protoplasts prepared from *N. tabacum* Bright Yellow-2 (BY-2) cells were performed as previously described (Vanden Bossche et al., 2013). Briefly, protoplasts were transfected with a *pC5-SD2::fLUC* reporter construct and effector constructs overexpressing *GUS*, *MYC1*, *MYC2*, *GAME9* or a combination thereof. A *pCaMV35S::rLUC* construct was co-transfected for normalization of fLUC activity. Two micrograms of each construct were transfected and total DNA added was equalized with a *pCaMV35S::GUS* control construct. After overnight incubation followed by lysis of the cells, the luciferase activities were measured using the Dual-Luciferase Reporter Assay System (Promega). Each assay was carried out in eight biological repeats. Statistical significance was determined by unpaired Student’s *t*-tests.

### Generation and Cultivation of Tomato Hairy Roots

*S. lycopersicum* (cultivar Moneymaker) seed sterilization, rhizogenic *Agrobacterium*-mediated transformation of tomato seedlings and cultivation of hairy roots were carried out as previously described (Harvey et al., 2008; Ron et al., 2014) with following modifications. Tomato seeds were surface-sterilized in 70% (v/v) ethanol for 5 min followed by 3% (v/v) NaOCl for 20 min and three washes with sterile water. Seeds were plated on Murashige and Skoog (MS) medium (pH 5.8) containing 4.3 g/L of MS (Duchefa), 0.5 g/L of MES, 10 g/L of sucrose, and 10 g/L of agar (Neogen) in Magenta boxes. Boxes were put in the dark at 4°C for two days, in the dark at 24°C for one day, and in a 24°C controlled photoperiodic growth chamber (16:8 photoperiods) for ca. two weeks until cotyledons were fully expanded and the true leaves were just emerged. Competent rhizogenic *Agrobacterium* (strain ATCC15834) cells were transformed by electroporation with the desired binary vector, plated on yeast extract broth (YEB) medium with 100 mg/L of spectinomycin, and incubated at 28°C for four days. Each transformed culture was inoculated from plate into liquid YEB medium with 100 mg/L of spectinomycin and incubated overnight at 28°C with shaking at 200 rpm. Each transformed culture was used to transform approximately 40 cotyledon explants. Using a scalpel, cotyledons were cut in half after cutting off their base and top, which was followed by immersion of the explants in *Agrobacterium* culture with an optical density of 0.2–0.3 at 600 nm in liquid MS medium for 20 min. Next, the explants were blotted on sterile Whatman filter paper and transferred with their adaxial side down to plates with MS medium (pH 5.8) containing 4.4 g/L of MS supplemented with vitamins (Duchefa), 0.5 g/L of MES, 30 g/L of sucrose, and 8 g/L of agar (Neogen) without antibiotics. After three to four days of incubation in the dark at 25°C (Oberpichler et al., 2008), the explants were transferred with their adaxial side down to plates with MS medium (pH 5.8) containing 4.4 g/L of MS supplemented with vitamins (Duchefa), 0.5 g/L of MES, 30 g/L of sucrose, and 8 g/L of agar (Neogen) with 200 mg/L of cefotaxime and 50 mg/L of kanamycin. These plates were returned to the dark at 25°C until hairy roots emerged from infected sites. Hairy roots were excised and cultured on fresh plates with MS medium (pH 5.8) containing 4.4 g/L of MS supplemented with vitamins (Duchefa), 0.5 g/L of MES, 30 g/L of sucrose, and 10 g/L of agar (Neogen) with 200 mg/L of cefotaxime and 50 mg/L of kanamycin. After three rounds of subculture on plates with MS medium of the same composition, hairy roots were subcultured every four weeks on plates with MS medium without antibiotics for maintenance.

### Identification of CRISPR-Cas9 Hairy Root Mutants

CRISPR-Cas9 mutants were identified as described previously (Swinnen et al., 2020). Genomic DNA was prepared from homogenized hairy root cultures using extraction buffer (pH 9.5) containing 0.1 M of tris(hydroxymethyl)aminomethane (Tris)-HCl, 0.25 M of KCl, and 0.01 M of ethylenediaminetetraacetic acid (EDTA). This mixture was incubated at 95°C for 10 min and subsequently cooled at 4°C for 5 min. After addition of 3% (w/v) BSA, collected supernatant was used as a template in a standard PCR reaction using GoTaq (Promega) with Cas9-specific primers or primers to amplify the gRNA(s) target region(s) (Supplemental Table 1). PCR amplicons containing the gRNA(s) target site(s) were purified using HighPrep PCR reagent (MAGBIO). After Sanger sequencing of the purified PCR amplicons with an amplification primer located approximately 200 bp from the Cas9 cleavage site, quantitative sequence trace data were decomposed using Inference of CRISPR Editing (ICE) CRISPR Analysis Tool (https://ice.synthego.com/#/).

### Gene Expression Analysis by Quantitative Real-Time PCR

Hairy roots were grown for eight days in liquid MS medium (pH 5.8) containing 4.4 g/L of MS supplemented with vitamins (Duchefa), 0.5 g/L of MES, and 30 g/L of sucrose. Three biological replicates per line were treated for 24 h with 50 μM of JA or an equal amount of ethanol by replacement of the medium. Hairy roots were rinsed with purified water, harvested by flash freezing in liquid nitrogen, and ground using the Mixer Mill 300 (Retch).

Messenger RNA was extracted from approximately 15 mg of homogenized tissue as reported previously (Townsley et al., 2015) with following modifications. Tissue was lysed using 500 µL of lysate binding buffer (LBB) containing 100 mM of Tris-HCl (pH 7.5), 500 mM of LiCl, 10 mM of EDTA (pH 8.0), 1% of sodium dodecyl sulfate (SDS), 5 mM of dithiothreitol (DTT), 15 μL/mL of Antifoam A, and 5 μL/mL of 2-mercaptoethanol, and the mixture was allowed to stand for 10 min. Messenger RNA was separated from 200 µL of lysate using 1 µL of 12.5 µM of 5’ biotinylated polyT oligonucleotide (5’-biotin-ACAGGACATTCGTCGCTTCCTTTTTTTTTTTTTTTTTTTT-3’) and the mixture was allowed to stand for 10 min. Next, captured messenger RNA was isolated from the lysate by adding 20 µL of LBB-washed streptavidin-coated magnetic beads (New England Biolabs) and was allowed to stand for 10 min. Samples were placed on a MagWell Magnetic Separator 96 (EdgeBio) and washed with 200 µL of washing buffer A (10 mM of Tris-HCl (pH 7.5), 150 mM of LiCl, 1 mM of EDTA (pH 8.0), 0.1% of SDS), washing buffer B (10 mM of Tris-HCl (pH 7.5), 150 mM of LiCl, 1 mM of EDTA (pH 8.0)), and low-salt buffer (20 mM of Tris-HCl (pH 7.5), 150 mM of NaCl, 1 mM of EDTA (pH 8.0)), which were pre-chilled on ice. Elution of messenger RNA was done by adding 20 µL of 10 mM of Tris-HCl (pH 8.0) with 1 mM of 2-mercaptoethanol followed by incubation of the mixture at 80°C for 2 min.

First-strand complementary DNA was synthesized from 20 µL of messenger RNA eluate by qScript cDNA Synthesis Kit (Quantabio). For control samples, cDNA of three biological replicates was pooled per independent line and treatment. Quantitative real-time PCR (qPCR) reactions were carried out with a LightCycler 480 System (Roche) using Fast SYBR Green Master Mix (Applied Biosystems) and primers (Supplemental Table 1) designed by QuantPrime (https://www.quantprime.de/) (Arvidsson et al., 2008). Gene expression levels were quantified relative to *CLATHRIN ADAPTOR COMPLEXES MEDIUM SUBUNIT* (*CAC*) and *TAP42-INTERACTING PROTEIN* (*TIP41*) using the 2^-ΔΔCt^ method (Livak and Schmittgen, 2001). Statistical significance of log_2_-transformed data was determined by ANOVA followed by Tukey post-hoc analysis (P < 0.05).

### Targeted Metabolite Profiling by Liquid Chromatography-Mass Spectrometry

Hairy roots were grown for four weeks in liquid MS medium (pH 5.8) containing 4.4 g/L of MS supplemented with vitamins (Duchefa), 0.5 g/L of MES, and 30 g/L of sucrose. Five biological replicates per line were treated for 24 h or 96 h with 50 μM of JA or an equal amount of ethanol by replacement of the medium. Hairy roots were rinsed with purified water, harvested by flash freezing in liquid nitrogen, and ground with pestle and mortar. Approximately 400 mg of homogenized tissue was extracted using 1 mL of MeOH at room temperature for 10 min. Next, supernatant was evaporated to dryness under vacuum, the residue was dissolved in 800 μL of H_2_O/cyclohexane (1:1, v/v), and 100 μL of the aqueous phase was filtered using a 0.2 μm filter plate (Pall) and retained for analysis.

For LC-MS, 10 µL of the sample was injected into an Acquity UPLC BEH C18 column (2.1 x 150 mm, 1.7 µm) mounted on a Waters Acquity UPLC system coupled to a SYNAPT HDMS Q-TOF via an electrospray ionization source operated negative mode. The following gradient was run using acidified (0.1% (v/v) formic acid) solvents A (water/acetonitrile, 99:1, v/v) and B (acetonitrile/water; 99:1, v/v) at a flow rate of 350 µL/min: time 0 min, 5% B; 30 min, 50% B; 33 min, 100% B. Negative mode MS and chromatogram integration and alignment using the Progenesis QI software package (Waters) were carried out as described (Vanholme et al., 2013). Statistical significance was determined by ANOVA followed by Tukey post-hoc analysis (P < 0.05) or by unpaired Student’s *t*-tests.

### Phylogenetic Analysis

Amino acid sequences from *A. thaliana* MYC2 orthologs were retrieved from the comparative genomics resource PLAZA 4.0 Dicots (http://bioinformatics.psb.ugent.be/plaza/) (Van Bel et al., 2018), with the exception of *N. tabacum* sequences that were retrieved through a BLASTP search in the National Center for Biotechnology Information (NCBI) GenBank protein database. All phylogenetic analyses were conducted in MEGA7 (Kumar et al., 2016). A multiple sequence alignment of the full-length proteins was generated with MUSCLE and can be found in Supplemental Figure 13. Using the Find Best DNA/Protein Models (ML) tool, the Maximum Likelihood method based on the Le_Gascuel_2008 model (Le and Gascuel, 2008) was chosen to infer the phylogenetic tree. Evolutionary rate differences among sites were modeled using a discrete Gamma distribution. All positions with less than 95% site coverage were eliminated. Bootstrap analysis was carried out with 1,000 replicates.

## Accession Numbers

Sequence data from this article can be found in the EMBL/GenBank/Solgenomics data libraries under the following accession numbers: *MYC2* (Solyc08g076930), *MYC1* (Solyc08g005050), *CAS* (Solyc04g070980), *3bHSD2* (Solyc02g081730), *CPI* (Solyc12g098640), *CYP51* (Solyc01g008110), *C14-R* (Solyc09g009040), *8,7-SI* (Solyc06g082980), *SSR2* (Solyc02g069490), *SMO3* (Solyc01g091320), *SMO4* (Solyc06g005750), *C5-SD2* (Solyc02g086180), *7-DR2* (Solyc06g074090), *GAME11* (Solyc07g043420), *GAME6* (Solyc07g043460), *GAME4* (Solyc12g006460), *GAME12* (Solyc12g006470), *GAME25* (Solyc01g073640), *GAME1* (Solyc07g043490), *GAME17* (Solyc07g043480), *GAME18* (Solyc07g043500), *GAME9* (Solyc01g090340), *CAC* (Solyc08g006960), *TIP41* (Solyc10g049850), *JAZ1* (Solyc12g009220), *ODC* (Solyc04g082030), and *COI1* (Solyc05g052620).

## ACKNOWLEDGMENTS

This work was supported by the Research Foundation Flanders (FWO) through the projects G005312N and G004515N, a predoctoral fellowship to E.C., postdoctoral fellowships to J.P., P.F.C., and L.P., and by the European Community’s Horizon2020 Program under grant agreement [760331-Newcotiana]. We thank Geert Goeminne (VIB Metabolomics Core) for processing of the LC-MS chromatograms and Annick Bleys for help with preparing the manuscript. In addition, we thank Siobhan Brady and Kaisa Kajala for training G.S. in the use of CRISPR-Cas9 genome editing and in the generation of tomato hairy roots while hosting her in Siobhan Brady’s lab.

## AUTHOR CONTRIBUTIONS

G.S., L.P., and A.G. designed the experiments. G.S., M.D.M., J.P., F.J.M.H., E.C., R.D.C., R.V.B., P.F.C., and M.R. performed experiments. G.S., J.P., L.P., and A.G. analyzed the data. G.S. wrote the article and J.P., L.P., and A.G. complemented the writing. A.G. agrees to serve as the author responsible for contact and ensures communication. All scientists who have contributed substantially to the conception, design or execution of the work described in the manuscript are included as authors, in accordance with the guidelines from the Committee on Publication Ethics (COPE) (http://publicationethics.org/resources/guidelines). All authors agree to the list of authors and the identified contributions.

## Supplemental Data

**Supplemental Figure 1.**
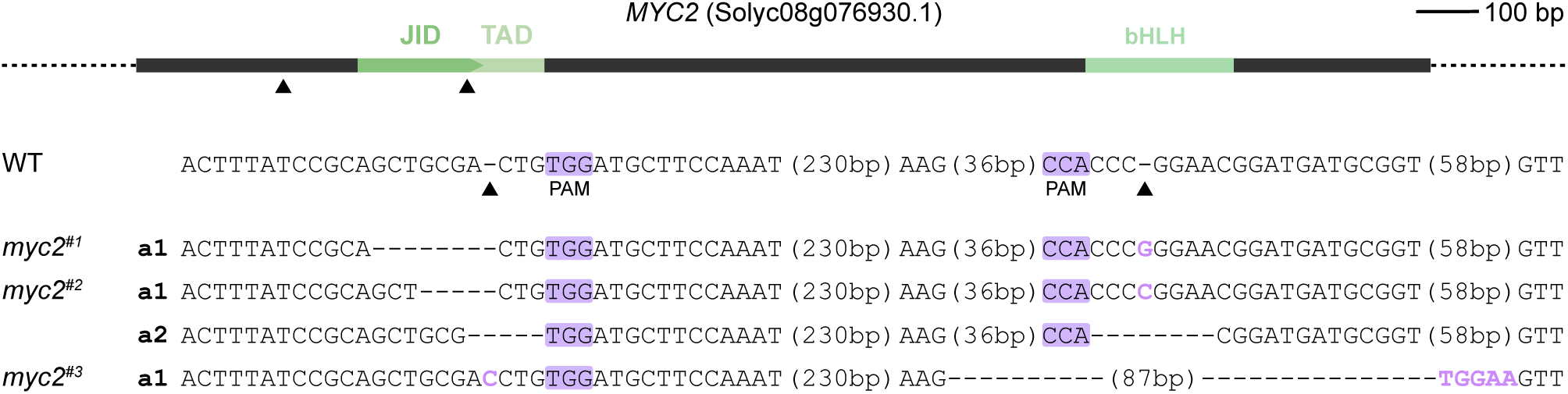
Schematic representation of *MYC2* with location of the CRISPR-Cas9 cleavage sites and *myc2* mutant sequences. The dark grey box represents the exon. Green boxes represent encoded protein domains. Cas9 cleavage sites for guide RNAs are indicated with arrowheads. Allele sequences of three independent *myc2* lines are shown. PAMs are marked in purple, inserted bases are shown in purple, deleted bases are replaced by a dash, and sequence gap length is shown between parentheses. Abbreviations: JID, JAZ interaction domain; TAD, transactivation domain; WT, wild-type; PAM, protospacer adjacent motif.

**Supplemental Figure 2.**
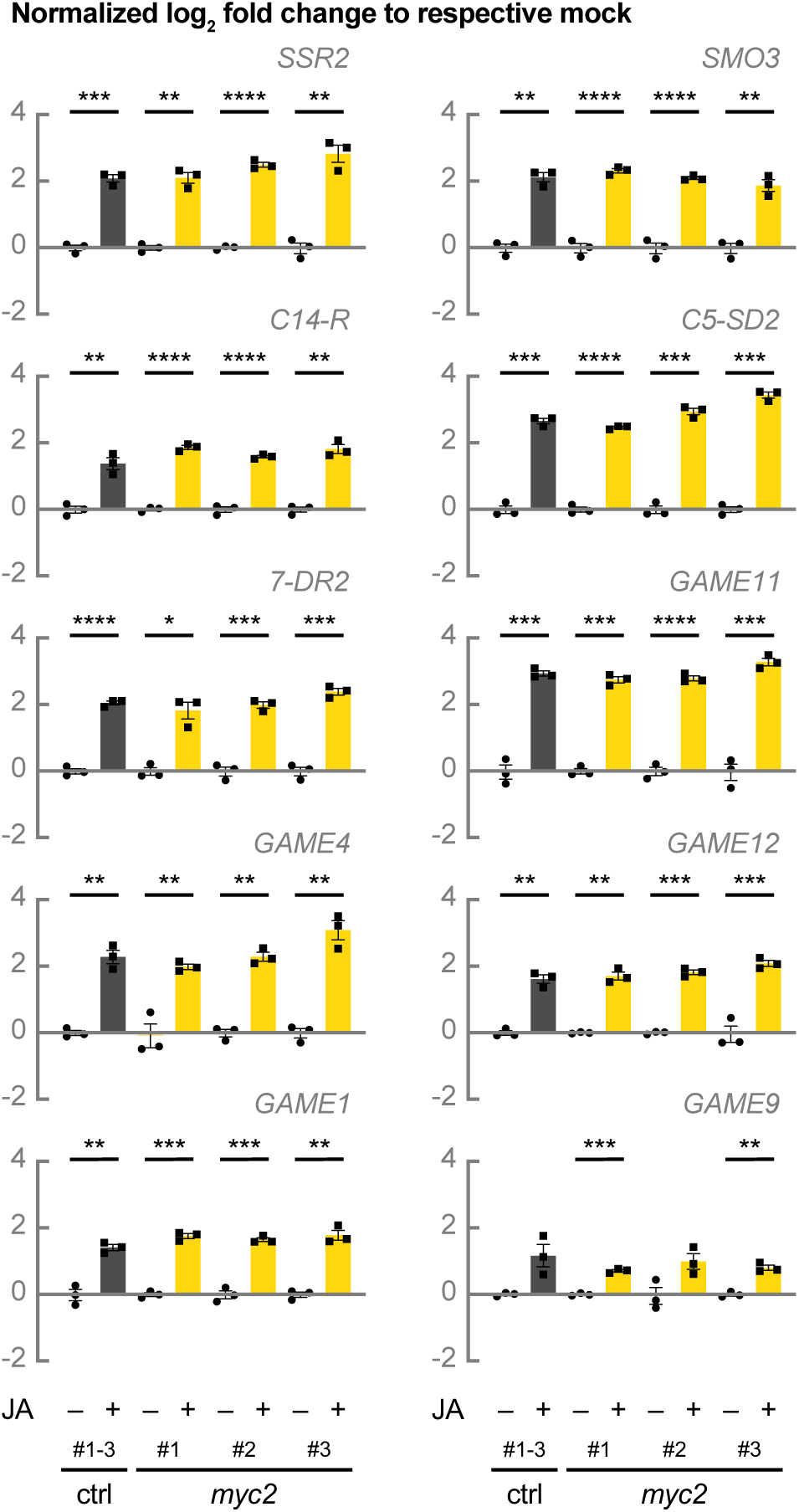
Unaffected JA inducibility of SGA biosynthesis gene expression in *myc2* lines. Alternative representation of relative expression of cholesterogenesis genes, SGA biosynthesis genes, and *GAME9* shown in Figure 1. Control hairy root lines expressing *pCaMV35S::GUS* (grey bars) and *myc2* lines (yellow bars) were treated for 24 h with 50 μM of JA or an equal amount of ethanol. For control samples, cDNA of three biological replicates was pooled per independent line and treatment. Bars represent mean log2-transformed fold changes relative to the mean of the respective mock-treated line. Error bars denote standard error (n = 3). Individual mock- (●) and JA-treated (■) values are shown. Statistical significance was determined by unpaired Student’s *t*-tests (*, P<0.05; **, P<0.01; ***, P<0.001; ****, P<0.0001). Abbreviations: *SSR2*, *STEROL SIDE CHAIN REDUCTASE 2*; *SMO3*, *C-4 STEROL METHYL OXIDASE 3*; *C14-R*, *STEROL C-14 REDUCTASE*; *7-DR2*, *7-DEHYDROCHOLESTEROL REDUCTASE 2*; *GUS*, *β-glucuronidase*.

**Supplemental Figure 3.**
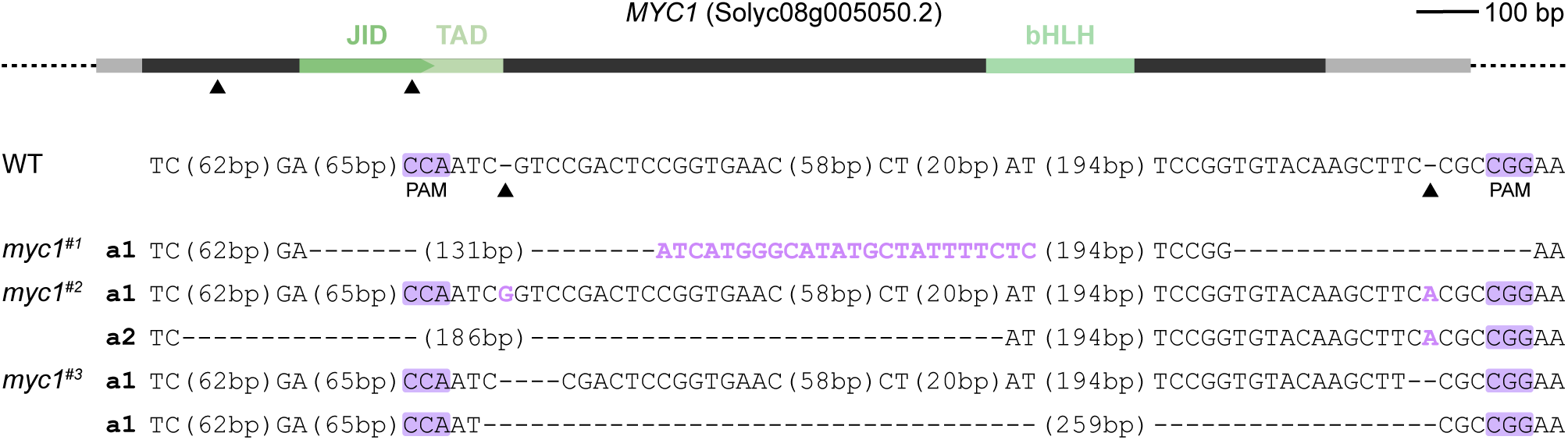
Schematic representation of *MYC1* with location of the CRISPR-Cas9 cleavage sites and *myc1* mutant sequences. The dark grey box represents the exon and light grey boxes represent UTRs. Green boxes represent encoded protein domains. Cas9 cleavage sites for guide RNAs are indicated with arrowheads. Allele sequences of three independent *myc1* lines are shown. PAMs are marked in purple, inserted bases are shown in purple, deleted bases are replaced by a dash, and sequence gap length is shown between parentheses. Abbreviations: JID, JAZ interaction domain; TAD, transactivation domain; WT, wild-type; PAM, protospacer adjacent motif; UTR, untranslated region.

**Supplemental Figure 4.**
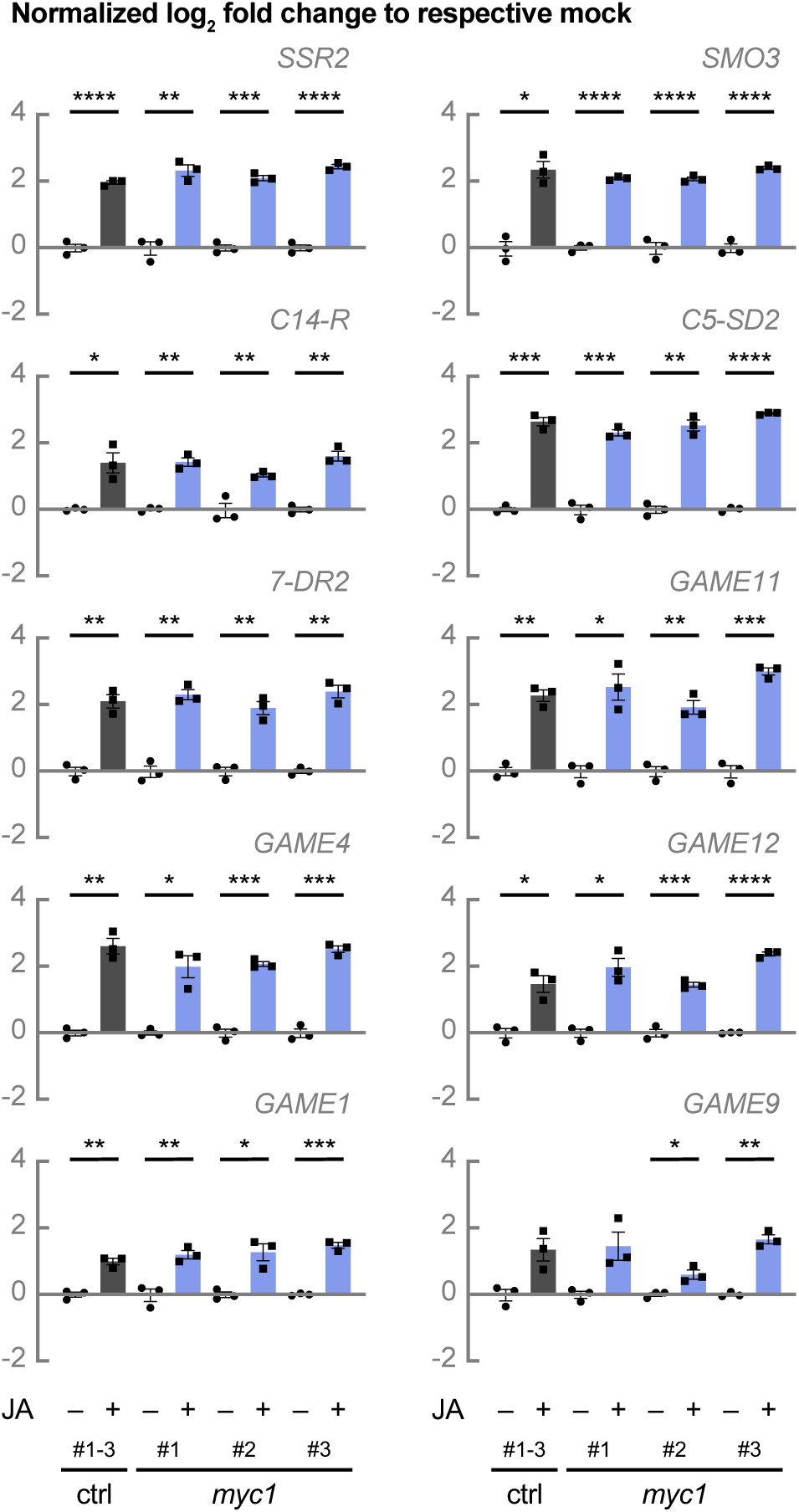
Unaffected JA inducibility of SGA biosynthesis gene expression in *myc1* lines. Alternative representation of relative expression of cholesterogenesis genes, SGA biosynthesis genes, and *GAME9* shown in Figure 5. Control hairy root lines expressing *pCaMV35S::GUS* (grey bars) and *myc1* lines (blue bars) were treated for 24 h with 50 μM of JA or an equal amount of ethanol. For control samples, cDNA of three biological replicates was pooled per independent line and treatment. Bars represent mean log2-transformed fold changes relative to the mean of the respective mock-treated line. Error bars denote standard error (n = 3). Individual mock- (●) and JA-treated (■) values are shown. Statistical significance was determined by unpaired Student’s *t*-tests (*, P<0.05; **, P<0.01; ***, P<0.001; ****, P<0.0001). Abbreviations: *SSR2*, *STEROL SIDE CHAIN REDUCTASE 2*; *SMO3*, *C-4 STEROL METHYL OXIDASE 3*; *C14-R*, *STEROL C-14 REDUCTASE*; *7-DR2*, *7-DEHYDROCHOLESTEROL REDUCTASE 2*; *GUS*, *β-glucuronidase*.

**Supplemental Figure 5.**
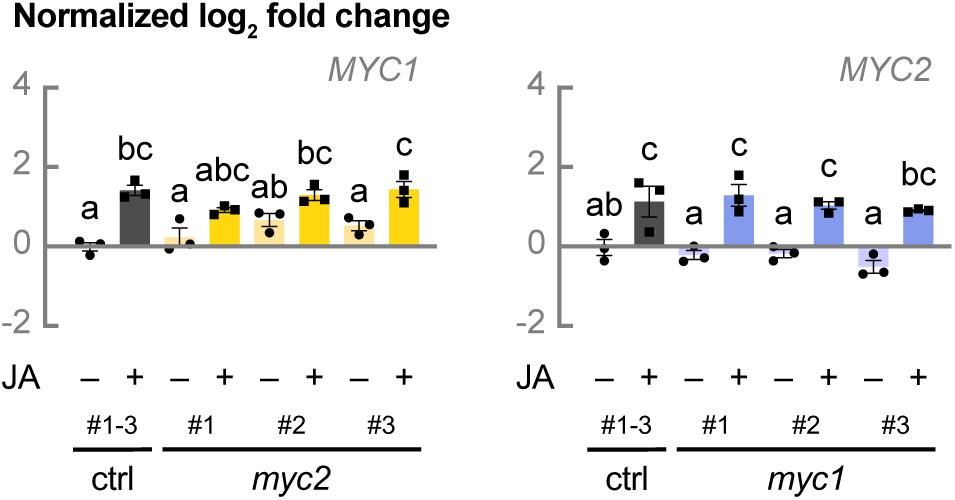
*myc2* and *myc1* lines do not exhibit upregulated expression of *MYC1* and *MYC2*, respectively. Relative expression of *MYC1* and *MYC2* analyzed by qPCR. Control hairy root lines expressing *pCaMV35S::GUS* (grey bars), *myc2* (yellow bars), and *myc1* lines (blue bars) were treated for 24 h with 50 μM of JA or an equal amount of ethanol. For control samples, cDNA of three biological replicates was pooled per independent line and treatment. Bars represent mean log_2_-transformed fold changes relative to the mean of three independent mock-treated control lines. Error bars denote standard error (n=3). Individual mock- (●) and JA-treated (■) values are shown. Statistical significance was determined by ANOVA followed by Tukey post-hoc analysis (P < 0.05; indicated by different letters). Abbreviations: *GUS*, *β-glucuronidase*.

**Supplemental Figure 6.**
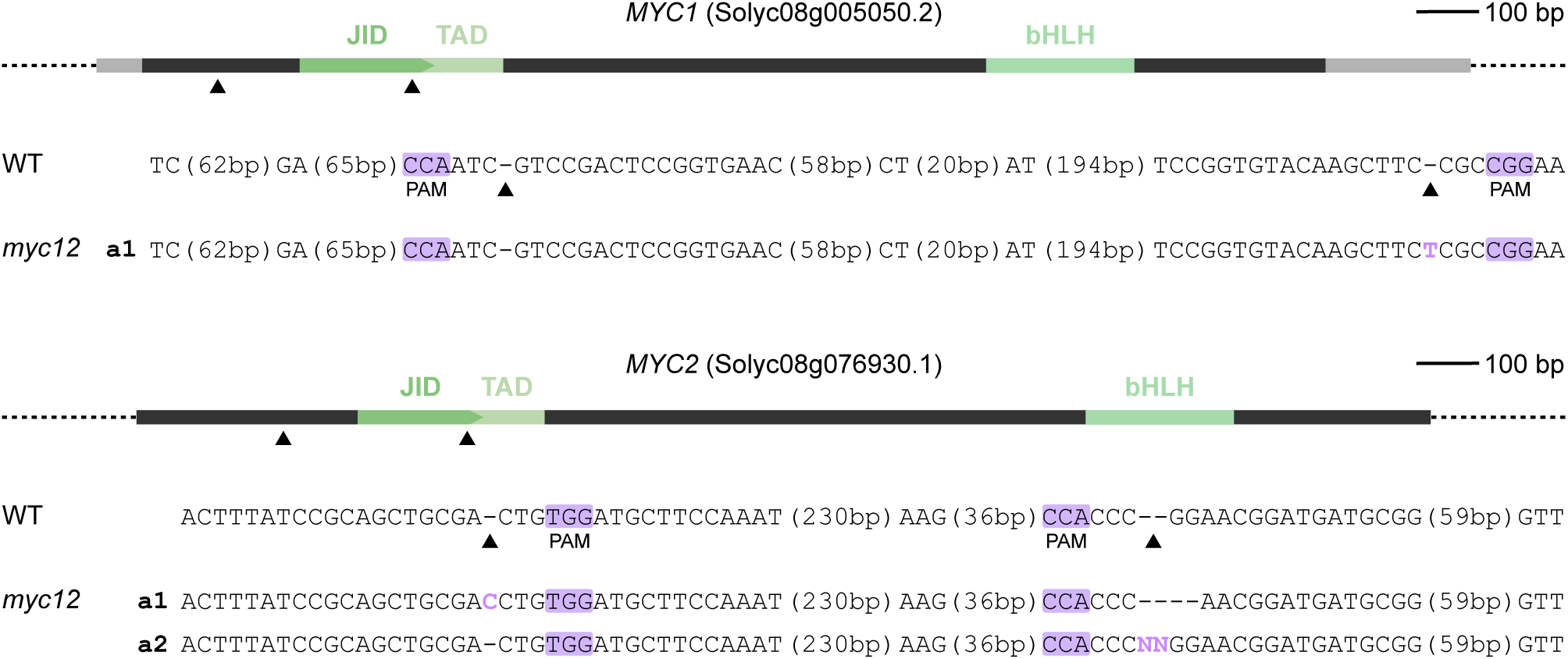
Schematic representation of *MYC1* and *MYC2* with location of the CRISPR-Cas9 cleavage sites and *myc1 myc2* mutant sequences. Dark grey boxes represent exons and light grey boxes represent UTRs. Green boxes represent encoded protein domains. Cas9 cleavage sites for guide RNAs are indicated with arrowheads. Allele sequences of one independent *myc1 myc2* (*myc12*) line are shown. PAMs are marked in purple, inserted bases are shown in purple, deleted bases are replaced by a dash, and sequence gap length is shown between parentheses. Abbreviations: JID, JAZ interaction domain; TAD, transactivation domain; WT, wild-type; PAM, protospacer adjacent motif; UTR, untranslated region.

**Supplemental Figure 7.**
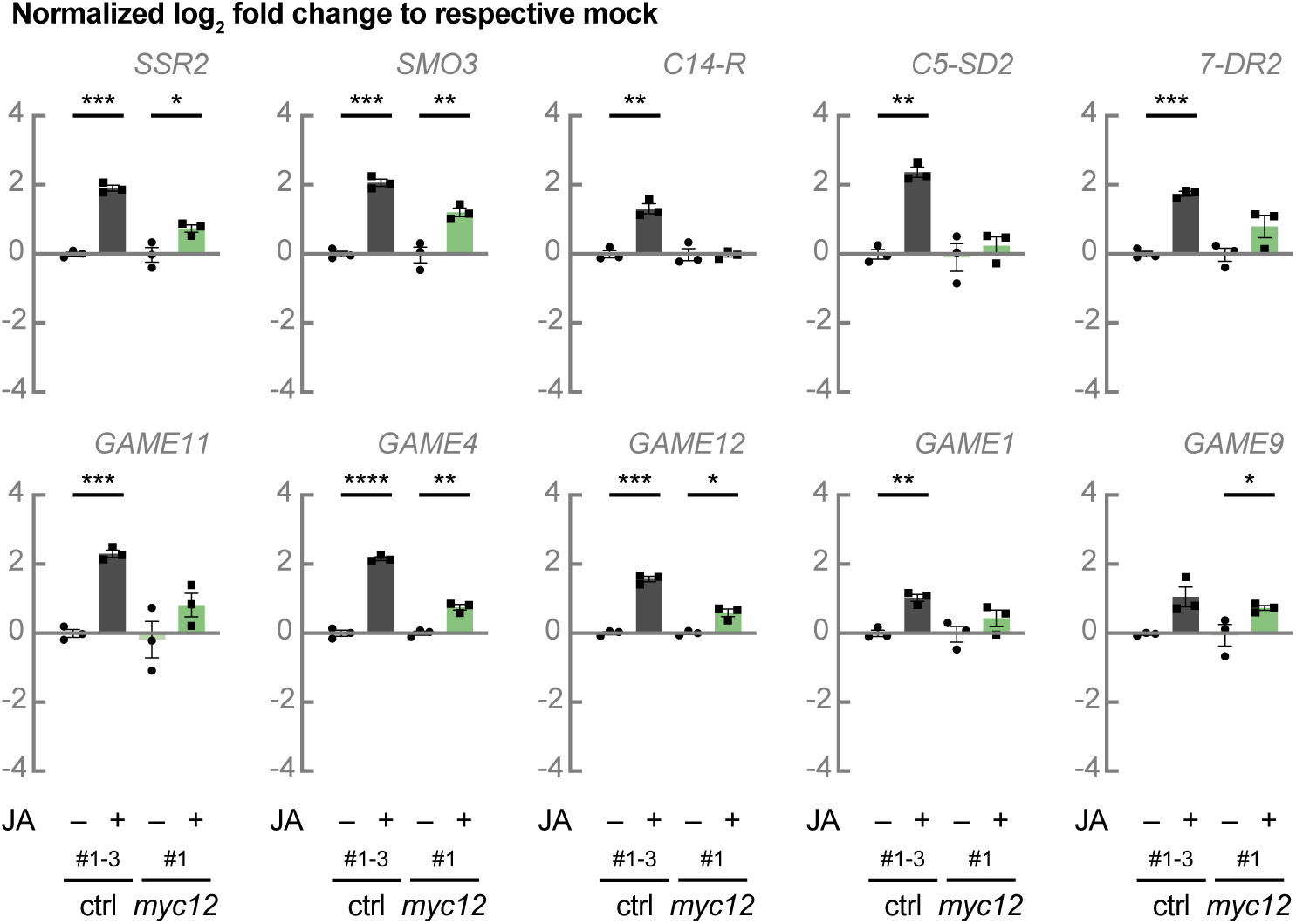
Reduced or absent JA inducibility of SGA biosynthesis gene expression in a *myc1 myc2* line. Alternative representation of relative expression of cholesterogenesis genes, SGA biosynthesis genes, and *GAME9* shown in Figure 7. Control hairy root lines expressing *pCaMV35S::GUS* (grey bars) and a *myc1 myc2* (*myc12*) line (green bars) were treated for 24 h with 50 μM of JA or an equal amount of ethanol. For control samples, cDNA of three biological replicates was pooled per independent line and treatment. Bars represent mean log_2_-transformed fold changes relative to the mean of the respective mock-treated line. Error bars denote standard error (n = 3). Individual mock- (●) and JA-treated (■) values are shown. Statistical significance was determined by unpaired Student’s *t*-tests (*, P<0.05; **, P<0.01; ***, P<0.001; ****, P<0.0001). Abbreviations: *SSR2*, *STEROL SIDE CHAIN REDUCTASE 2*; *SMO3*, *C-4 STEROL METHYL OXIDASE 3*; *C14-R*, *STEROL C-14 REDUCTASE*; *7-DR2*, *7-DEHYDROCHOLESTEROL REDUCTASE 2*; *GUS*, *β-glucuronidase*.

**Supplemental Figure 8.**
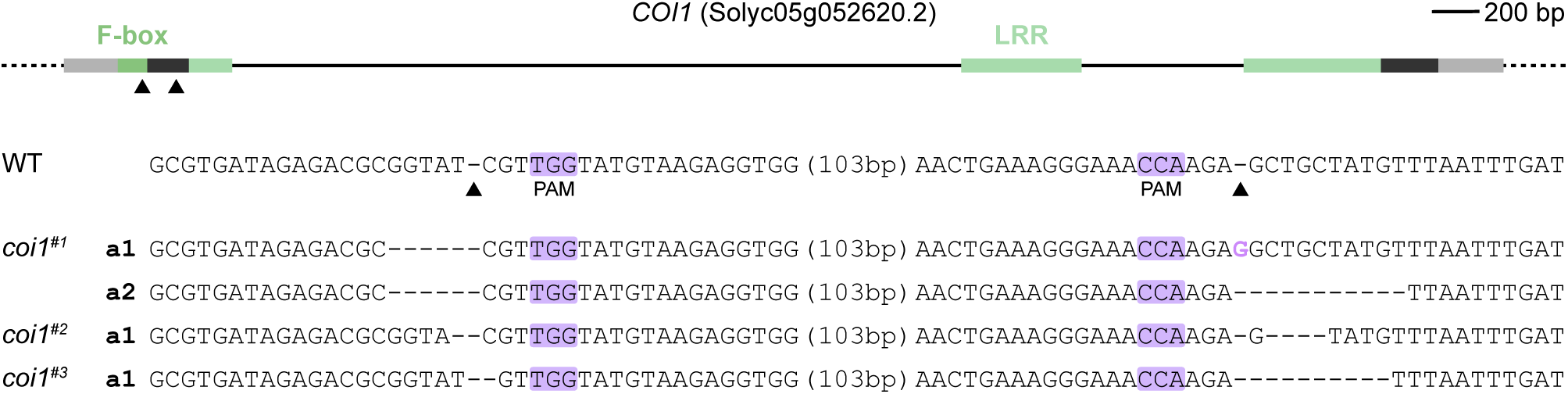
Schematic representation of *COI1* with location of the CRISPR-Cas9 cleavage sites and *coi1* mutant sequences. Dark grey boxes represent exons, solid lines introns, and light grey boxes UTRs. Green boxes represent encoded protein domains. Cas9 cleavage sites for guide RNAs are indicated with arrowheads. Allele sequences of three independent *coi1* lines are shown. PAMs are marked in purple, inserted bases are shown in purple, deleted bases are replaced by a dash, and sequence gap length is shown between parentheses. Abbreviations: LRR, leucine-rich repeat; WT, wild-type; PAM, protospacer adjacent motif; UTR, untranslated region.

**Supplemental Figure 9.**
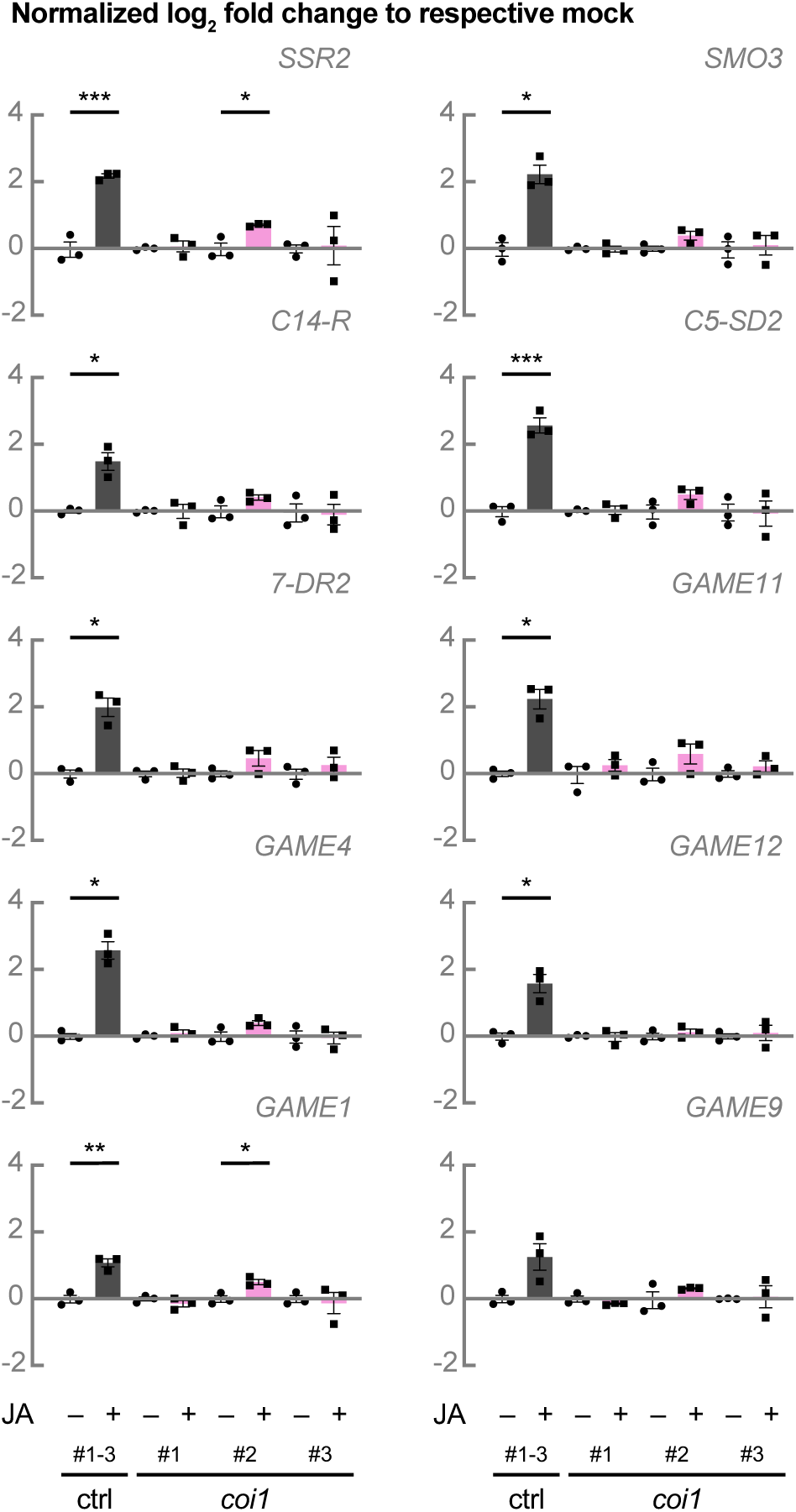
SGA biosynthesis gene expression is no longer JA inducible in *coi1* lines. Alternative representation of relative expression of cholesterogenesis genes, SGA biosynthesis genes, and *GAME9* shown in Figure 9. Control hairy root lines expressing *pCaMV35S::GUS* (grey bars) and *coi1* lines (pink bars) were treated for 24 h with 50 μM of JA or an equal amount of ethanol. For control samples, cDNA of three biological replicates was pooled per independent line and treatment. Bars represent mean log2-transformed fold changes relative to the mean of the respective mock-treated line. Error bars denote standard error (n = 3). Individual mock- (●) and JA-treated (■) values are shown. Statistical significance was determined by unpaired Student’s *t*-tests (*, P<0.05; **, P<0.01; ***, P<0.001; ****, P<0.0001). Abbreviations: *SSR2*, *STEROL SIDE CHAIN REDUCTASE 2*; *SMO3*, *C-4 STEROL METHYL OXIDASE 3*; *C14-R*, *STEROL C-14 REDUCTASE*; *7-DR2*, *7-DEHYDROCHOLESTEROL REDUCTASE 2*; *GUS*, *β-glucuronidase*.

**Supplemental Figure 10.**
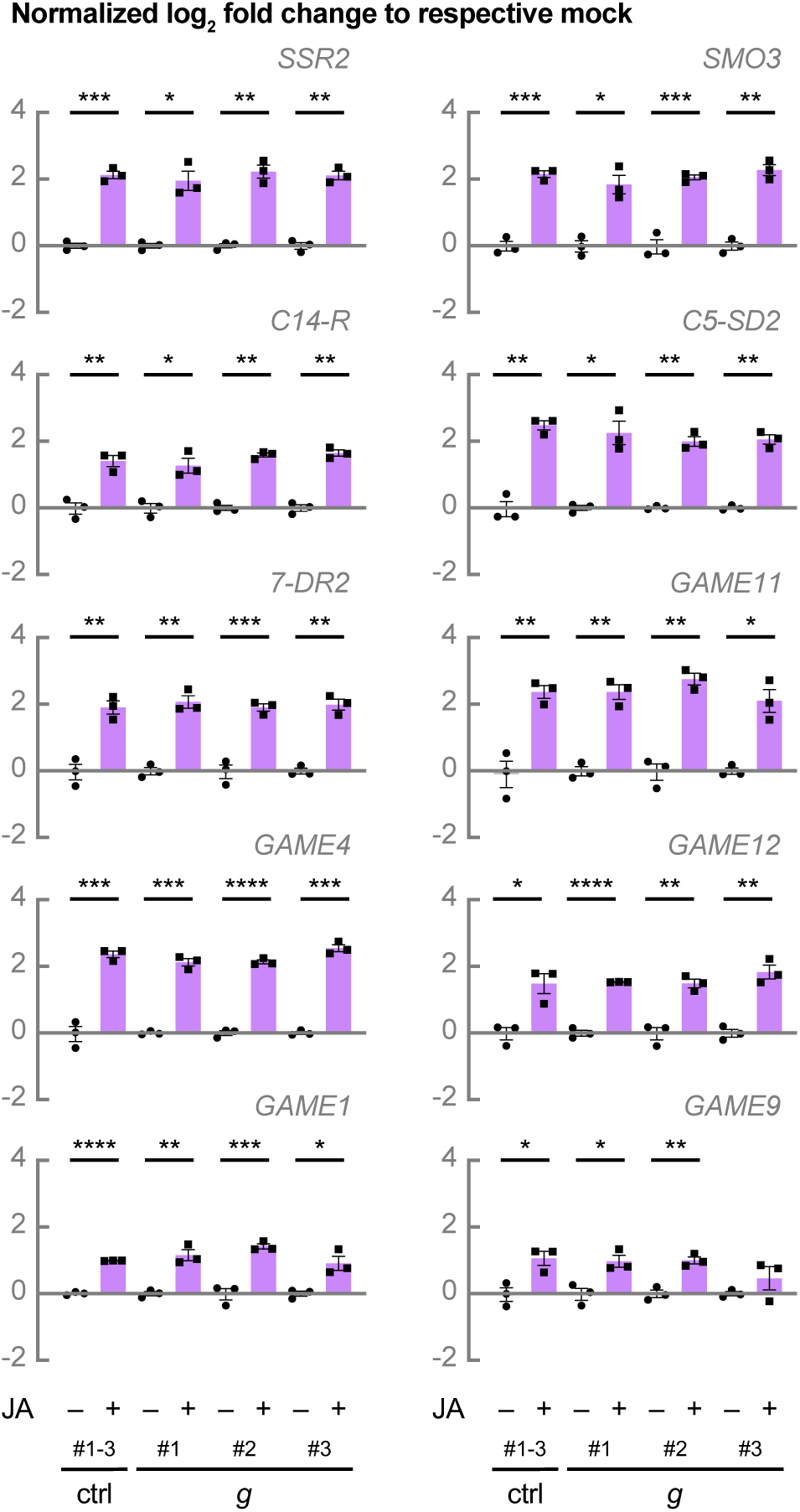
JA inducibility of *C5-SD2* expression is not affected in G-box (*g*) mutant lines. Alternative representation of relative expression of cholesterogenesis genes, SGA biosynthesis genes, and *GAME9* shown in Figure 11. Control hairy root lines expressing *pCaMV35S::GUS* (grey bars) and *g* lines (purple bars) were treated for 24 h with 50 μM of JA or an equal amount of ethanol. For control samples, cDNA of three biological replicates was pooled per independent line and treatment. Bars represent mean log2-transformed fold changes relative to the mean of the respective mock-treated line. Error bars denote standard error (n = 3). Individual mock- (●) and JA-treated (■) values are shown. Statistical significance was determined by unpaired Student’s *t*-tests (*, P<0.05; **, P<0.01; ***, P<0.001; ****, P<0.0001). Abbreviations: *SSR2*, *STEROL SIDE CHAIN REDUCTASE 2*; *SMO3*, *C-4 STEROL METHYL OXIDASE 3*; *C14-R*, *STEROL C-14 REDUCTASE*; *7-DR2*, *7-DEHYDROCHOLESTEROL REDUCTASE 2*; *GUS*, *β-glucuronidase*.

**Supplemental Figure 11.**
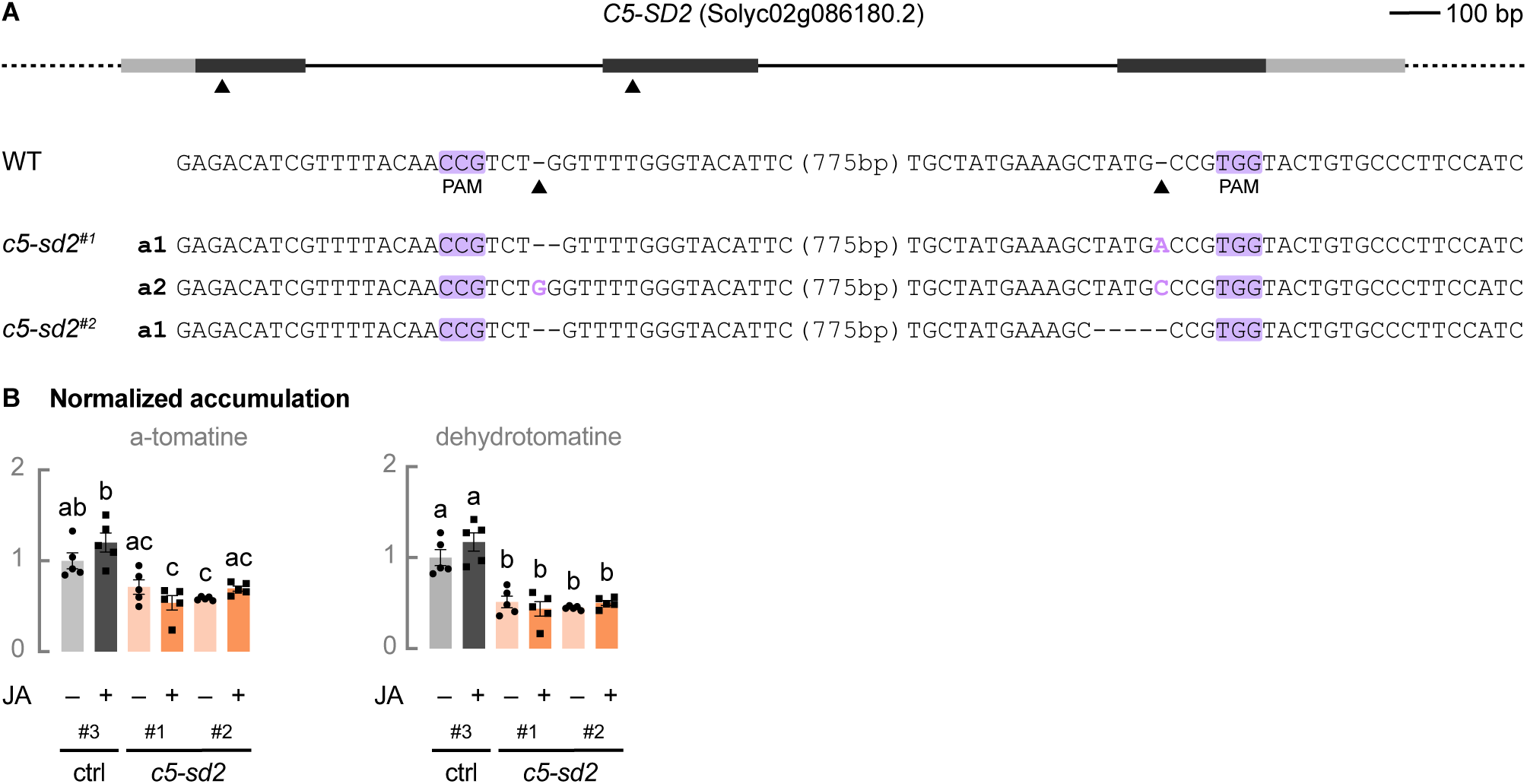
Reduced SGA levels in *c5-sd2* lines. (A) Schematic representation of *C5-SD2* with location of the CRISPR-Cas9 cleavage sites and *c5-sd2* mutant sequences. Dark grey boxes represent exons, solid lines introns, and light grey boxes UTRs. Cas9 cleavage sites for guide RNAs are indicated with arrowheads. Allele sequences of two independent *c5-sd2* lines are shown. PAMs are marked in purple, inserted bases are shown in purple, deleted bases are replaced by a dash, and sequence gap length is shown between parentheses. (B) Relative accumulation of *α*-tomatine and dehydrotomatine analyzed by LC-MS. Control hairy root lines expressing *pCaMV35S::GUS* (grey bars) and *c5-sd2* lines (orange bars) were treated for 24 h with 50 μM of JA or an equal amount of ethanol. Bars represent mean fold changes relative to the mean of the mock-treated control. Error bars denote standard error (n=5). Individual mock- (●) and JA-treated (■) values are shown. Statistical significance was determined by ANOVA followed by Tukey post-hoc analysis (P<0.05; indicated by different letters). Abbreviations: WT, wild-type; PAM, protospacer adjacent motif; UTR, untranslated region; *GUS*, *β-glucuronidase*.

**Supplemental Figure 12.**
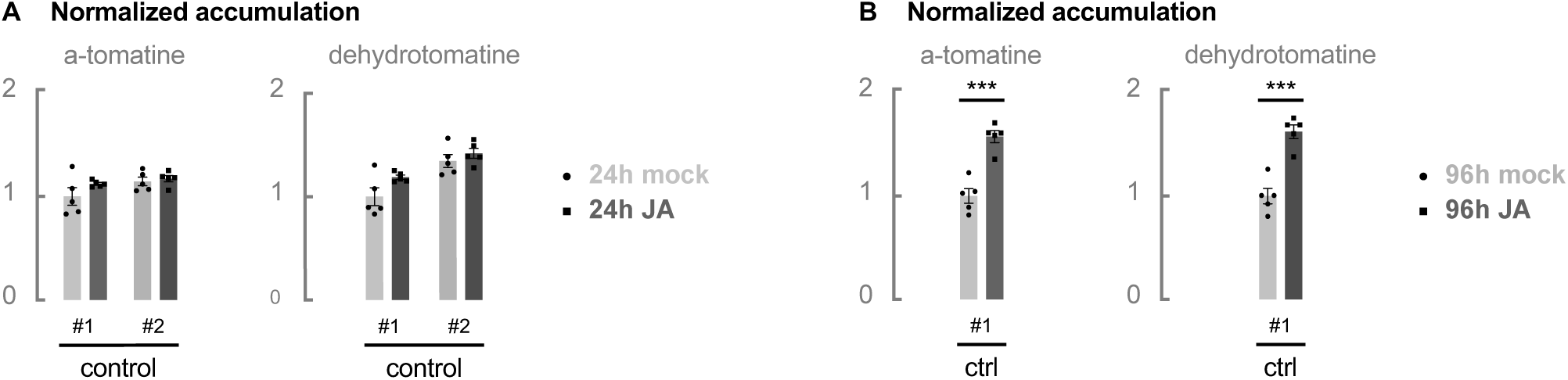
Limited SGA induction upon 96 h of JA treatment. Relative accumulation of *α*-tomatine and dehydrotomatine analyzed by LC-MS. Control hairy root lines expressing *pCaMV35S::GUS* were treated for 24 h (A) or 96 h (B) with 50 μM of JA or an equal amount of ethanol. Bars represent mean fold changes relative to the mean of the leftmost mock-treated control. Error bars denote standard error (n=5). Individual mock- (●) and JA-treated (■) values are shown. Statistical significance was determined by unpaired Student’s *t*-tests (*, P<0.05; **, P<0.01; ***, P<0.001; ****, P<0.0001). Abbreviations: *GUS*, *β-glucuronidase*.

**Supplemental Figure 13.**
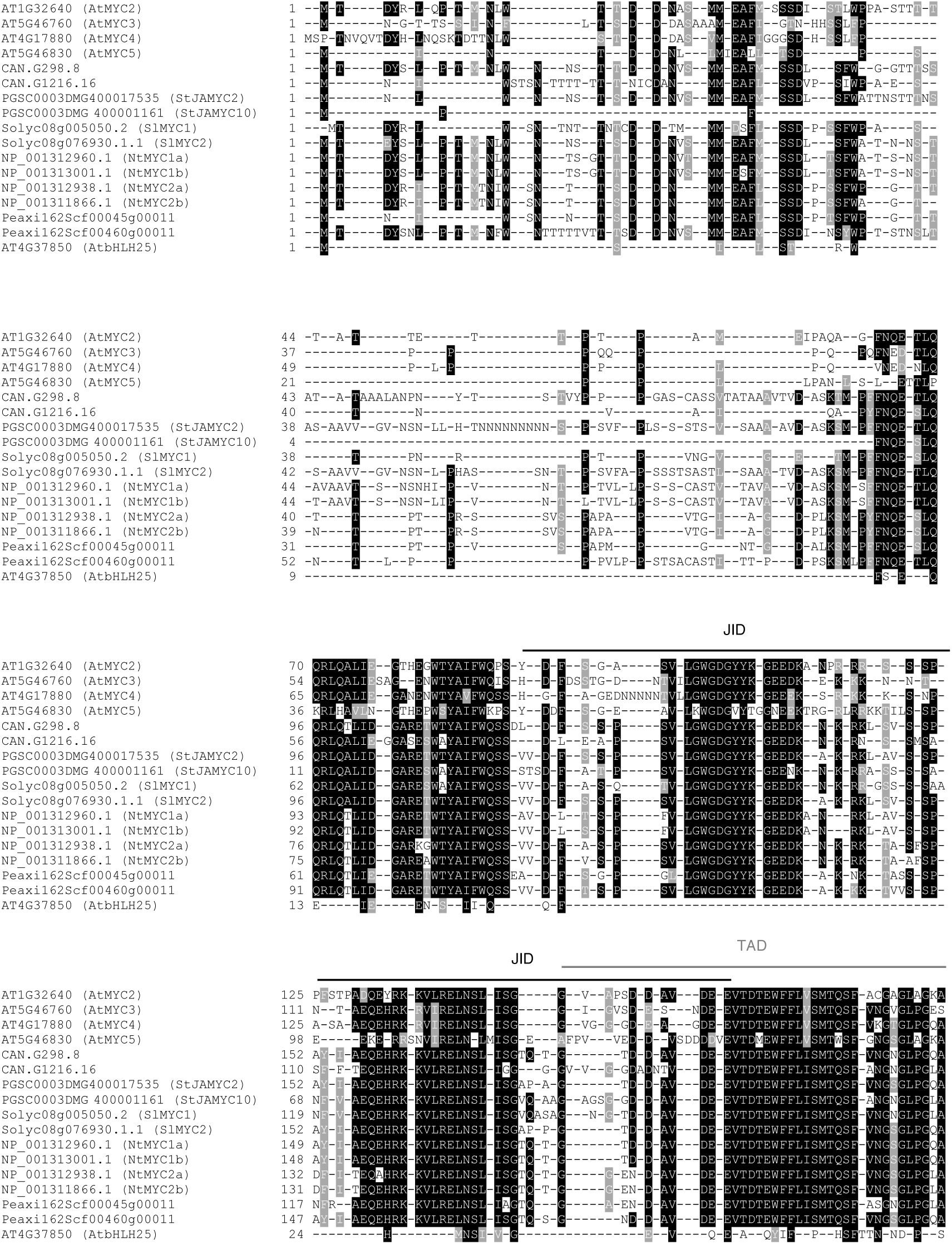
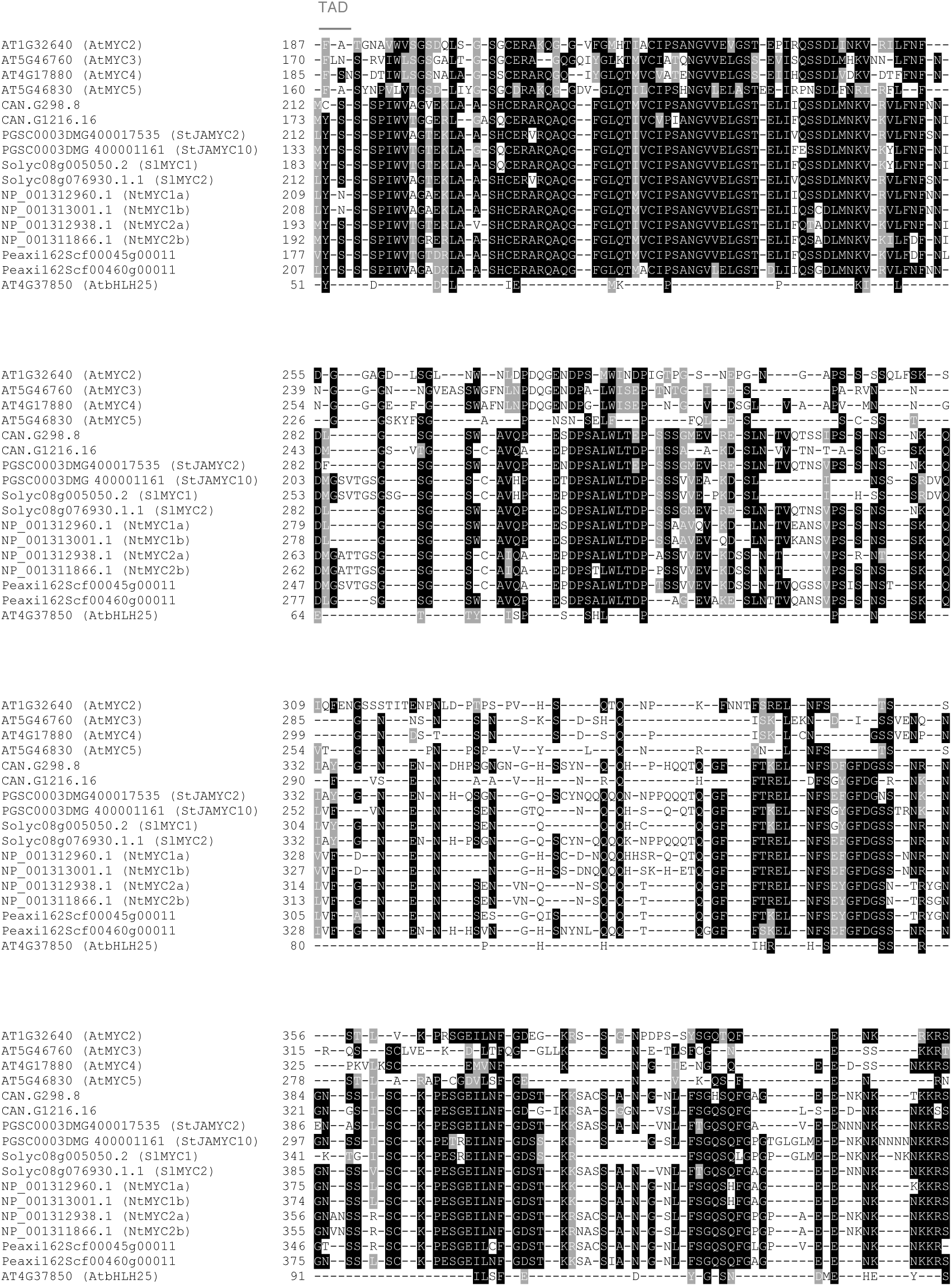
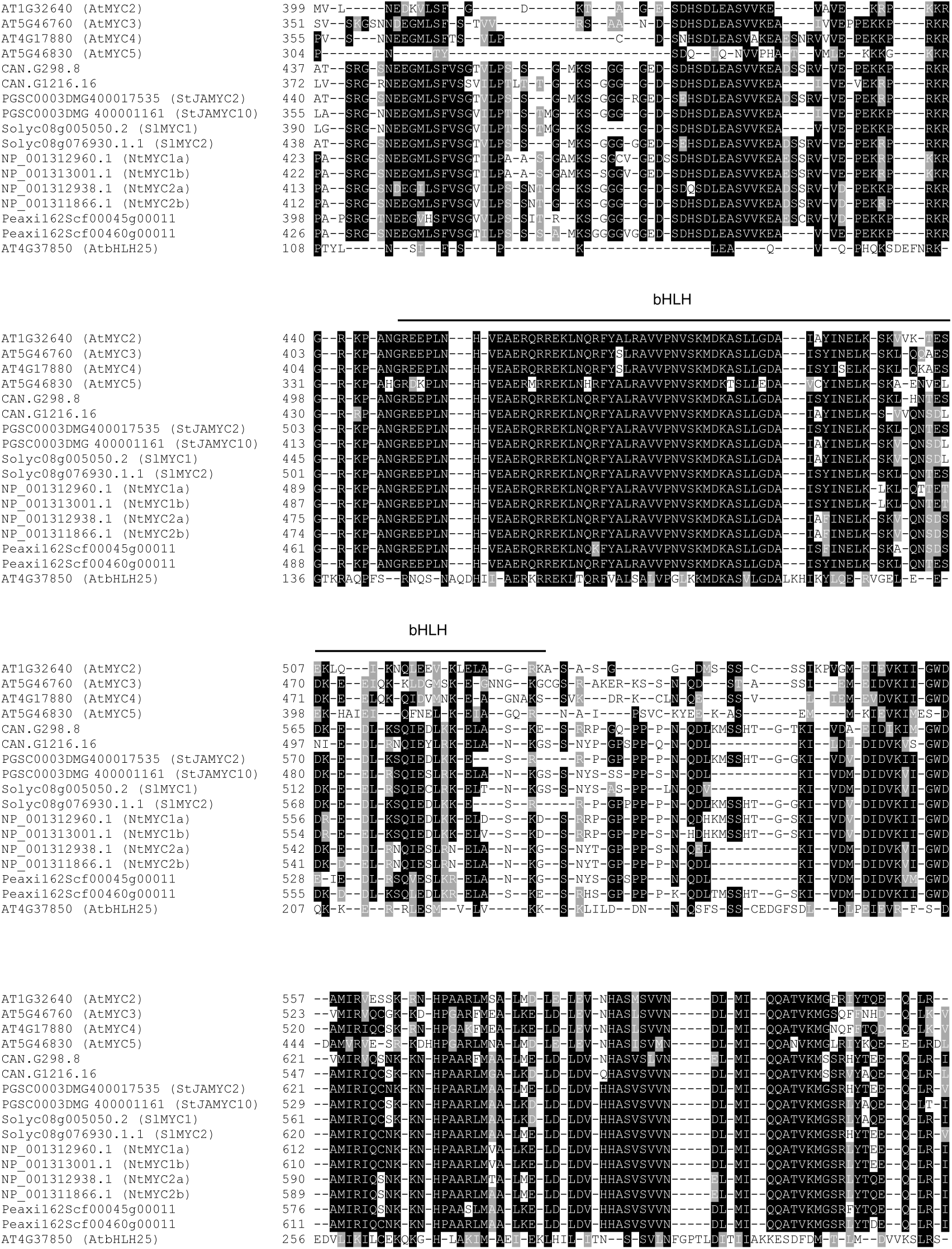
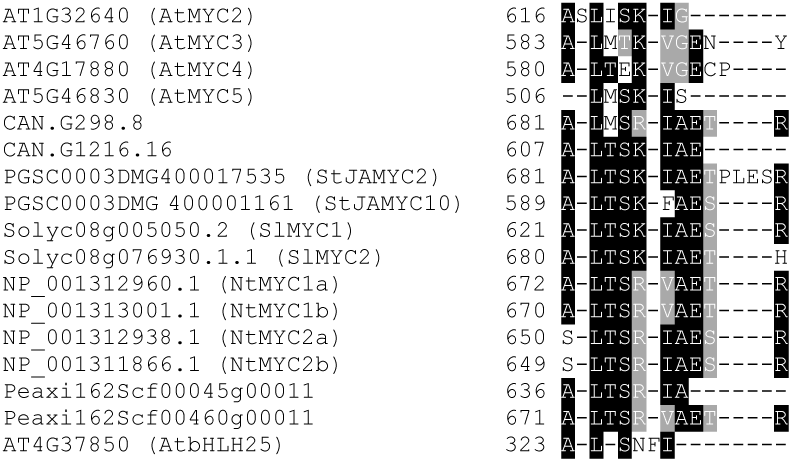
Protein alignment generated by MUSCLE used for the phylogenetic tree in Figure 3A.

**Supplemental Table 1.**
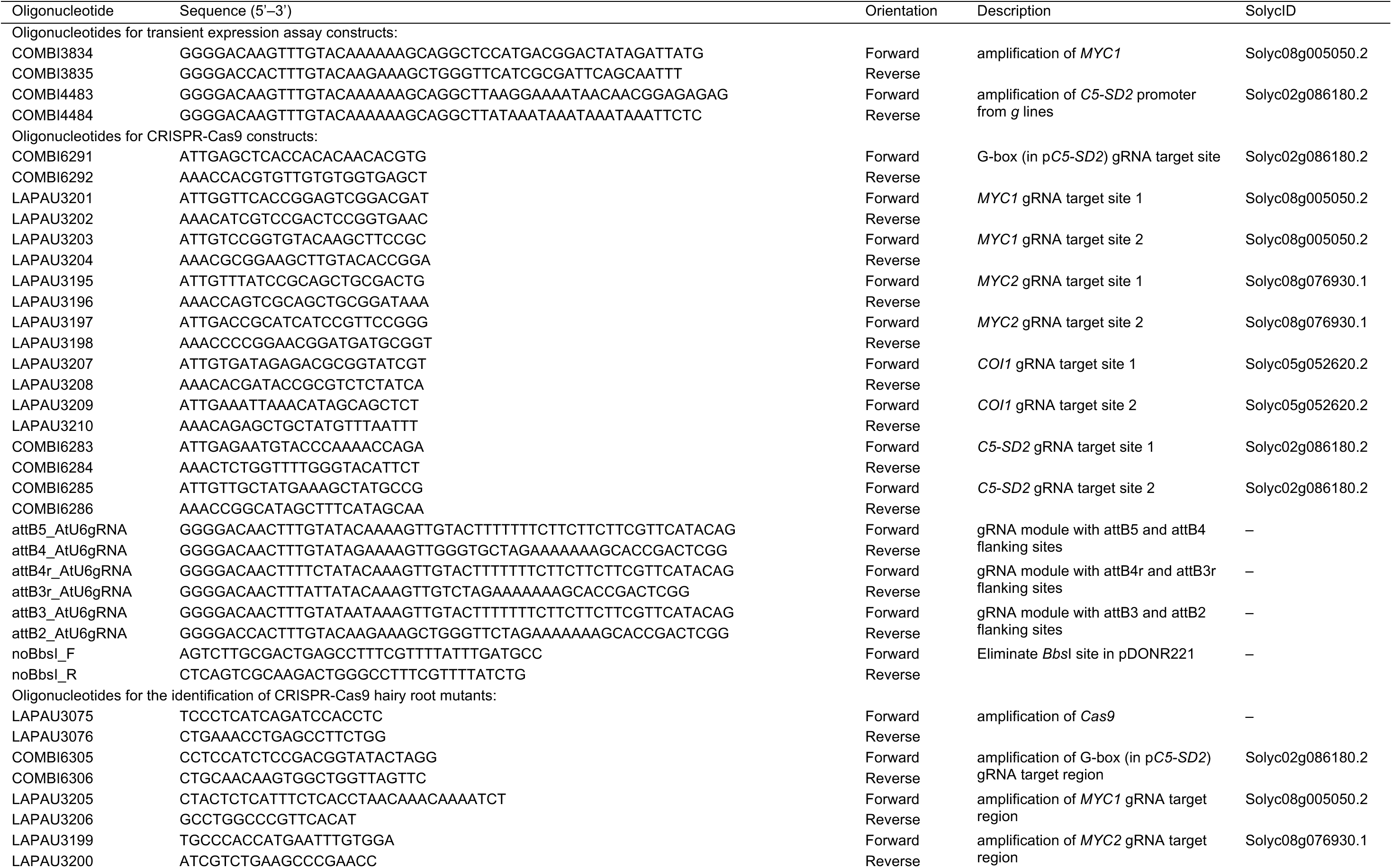
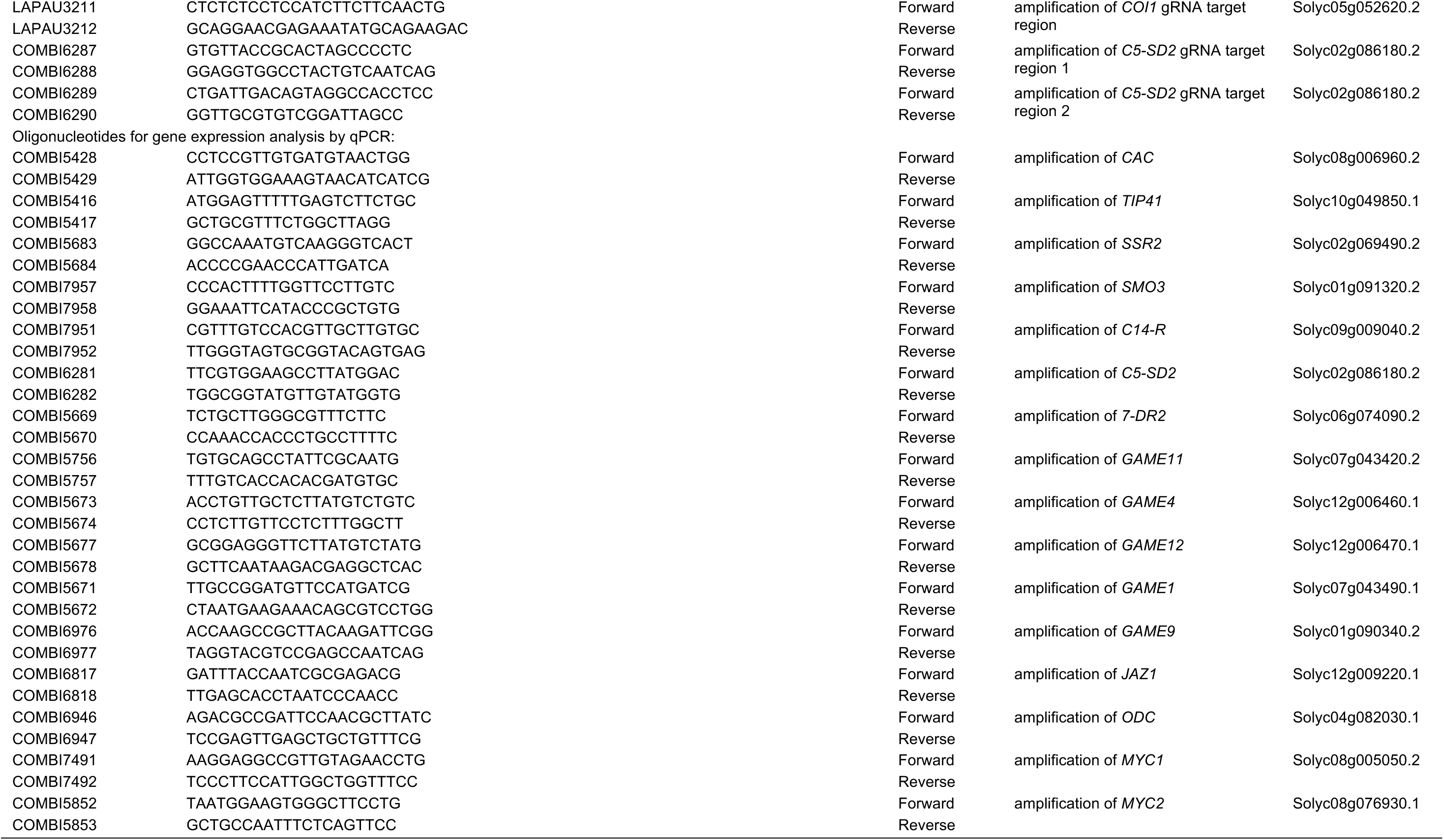
Oligonucleotides used in this study

## REFERENCES

Abdelkareem, A., Thagun, C., Nakayasu, M., Mizutani, M., Hashimoto, T., and Shoji, T. (2017). Jasmonate-induced biosynthesis of steroidal glycoalkaloids depends on COI1 proteins in tomato. Biochem. Biophys. Res. Commun. 489: 206–210.

Acosta, C., Pérez-Amador, M.A., Carbonell, J., and Granell, A. (2005). The two ways to produce putrescine in tomato are cell-specific during normal development. Plant Sci. 168: 1053–1057.

Arvidsson, S., Kwasniewski, M., Riaño-Pachón, D.M., and Mueller-Roeber, B. (2008). QuantPrime - a flexible tool for reliable high-throughput primer design for quantitative PCR. BMC Bioinformatics 9: 465.

Boter, M., Ruíz-Rivero, O., Abdeen, A., and Prat, S. (2004). Conserved MYC transcription factors play a key role in jasmonate signaling both in tomato and *Arabidopsis*. Genes Dev. 18: 1577–1591.

Brkljacic, J., and Grotewold, E. (2017). Combinatorial control of plant gene expression. Biochim. Biophys. Acta - Gene Regul. Mech. 1860: 31–40.

Cárdenas, P.D., et al. (2016). GAME9 regulates the biosynthesis of steroidal alkaloids and upstream isoprenoids in the plant mevalonate pathway. Nat. Commun. 7: 10654.

Cárdenas, P.D., et al. (2019). Pathways to defense metabolites and evading fruit bitterness in genus *Solanum* evolved through 2-oxoglutarate-dependent dioxygenases. Nat. Commun. 10: 5169.

Chen, H., Jones, A.D., and Howe, G.A. (2006). Constitutive activation of the jasmonate signaling pathway enhances the production of secondary metabolites in tomato. FEBS Lett. 580: 2540–2546.

Chini, A., Gimenez-Ibanez, S., Goossens, A., and Solano, R. (2016). Redundancy and specificity in jasmonate signalling. Curr. Opin. Plant Biol. 33: 147–156.

Chini, A., Ben-Romdhane, W., Hassairi, A., and Aboul-Soud, M.A.M. (2017). Identification of TIFY/JAZ family genes in *Solanum lycopersicum* and their regulation in response to abiotic stresses. PLoS ONE 12: e0177381.

Chini, A., Fonseca, S., Fernández, G., Adie, B., Chico, J.M., Lorenzo, O., García-Casado, G., López-Vidriero, I., Lozano, F.M., Ponce, M.R., Micol, J.L., and Solano, R. (2007). The JAZ family of repressors is the missing link in jasmonate signalling. Nature 448: 666–671.

De Geyter, N., Gholami, A., Goormachtig, S., and Goossens, A. (2012). Transcriptional machineries in jasmonate-elicited plant secondary metabolism. Trends Plant Sci. 17: 349–359.

Du, M., et al. (2017). MYC2 orchestrates a hierarchical transcriptional cascade that regulates jasmonate-mediated plant immunity in tomato. Plant Cell 29: 1883–1906.

Fauser, F., Schiml, S., and Puchta, H. (2014). Both CRISPR/Cas-based nucleases and nickases can be used efficiently for genome engineering in *Arabidopsis thaliana*. Plant J. 79: 348–359.

Fonseca, S., Chini, A., Hamberg, M., Adie, B., Porzel, A., Kramell, R., Miersch, O., Wasternack, C., and Solano, R. (2009). (+)-7-*iso*-Jasmonoyl-L-isoleucine is the endogenous bioactive jasmonate. Nat. Chem. Biol. 5: 344–350.

Friedman, M. (2002). Tomato glycoalkaloids: role in the plant and in the diet. J. Agric. Food Chem. 50: 5751–5780.

Friedman, M., and Levin, C.E. (1998). Dehydrotomatine content in tomatoes. J. Agric. Food Chem. 46: 4571–4576.

Goossens, J., Mertens, J., and Goossens, A. (2017). Role and functioning of bHLH transcription factors in jasmonate signalling. J. Exp. Bot. 68: 1333–1347.

Haeussler, M., Schönig, K., Eckert, H., Eschstruth, A., Mianné, J., Renaud, J.-B., Schneider-Maunoury, S., Shkumatava, A., Teboul, L., Kent, J., Joly, J.-S., and Concordet, J.-P. (2016). Evaluation of off-target and on-target scoring algorithms and integration into the guide RNA selection tool CRISPOR. Genome Biol. 17: 148.

Harvey, J.J.W., Lincoln, J.E., and Gilchrist, D.G. (2008). Programmed cell death suppression in transformed plant tissue by tomato cDNAs identified from an *Agrobacterium rhizogenes*-based functional screen. Mol. Genet. Genomics 279: 509–521.

Itkin, M., et al. (2011). GLYCOALKALOID METABOLISM1 is required for steroidal alkaloid glycosylation and prevention of phytotoxicity in tomato. Plant Cell 23: 4507–4525.

Itkin, M., et al. (2013). Biosynthesis of antinutritional alkaloids in solanaceous crops is mediated by clustered genes. Science 341: 175–179.

Kajikawa, M., Sierro, N., Kawaguchi, H., Bakaher, N., Ivanov, N.V., Hashimoto, T., and Shoji, T. (2017). Genomic insights into the evolution of the nicotine biosynthesis pathway in tobacco. Plant Physiol. 174: 999–1011.

Kazan, K., and Manners, J.M. (2013). MYC2: the master in action. Mol. Plant 6: 686–703.

Kleinstiver, B.P., Prew, M.S., Tsai, S.Q., Topkar, V.V., Nguyen, N.T., Zheng, Z., Gonzales, A.P.W., Li, Z., Peterson, R.T., Yeh, J.-R.J., Aryee, M.J., and Joung, J.K. (2015). Engineered CRISPR-Cas9 nucleases with altered PAM specificities. Nature 523: 481–485.

Kozukue, N., Han, J.-S., Lee, K.-R., and Friedman, M. (2004). Dehydrotomatine and α-tomatine content in tomato fruits and vegetative plant tissues. J. Agric. Food Chem. 52: 2079–2083.

Kumar, S., Stecher, G., and Tamura, K. (2016). MEGA7: Molecular Evolutionary Genetics Analysis version 7.0 for bigger datasets. Mol. Biol. Evol. 33: 1870–1874.

Le, S.Q., and Gascuel, O. (2008). An improved general amino acid replacement matrix. Mol. Biol. Evol. 25: 1307–1320.

Livak, K.J., and Schmittgen, T.D. (2001). Analysis of relative gene expression data using real-time quantitative PCR and the 2^-ΔΔCT^ method. Methods 25: 402–408.

Ma, Z., Zhu, P., Shi, H., Guo, L., Zhang, Q., Chen, Y., Chen, S., Zhang, Z., Peng, J., and Chen, J. (2019). PTC-bearing mRNA elicits a genetic compensation response via Upf3a and COMPASS components. Nature 568: 259–263.

Mertens, J., Van Moerkercke, A., Vanden Bossche, R., Pollier, J., and Goossens, A. (2016). Clade IVa basic helix-loop-helix transcription factors form part of a conserved jasmonate signaling circuit for the regulation of bioactive plant terpenoid biosynthesis. Plant Cell Physiol. 57: 2564–2575.

Montero-Vargas, J.M., Casarrubias-Castillo, K., Martínez-Gallardo, N., Ordaz-Ortiz, J.J., Délano-Frier, J.P., and Winkler, R. (2018). Modulation of steroidal glycoalkaloid biosynthesis in tomato (*Solanum lycopersicum*) by jasmonic acid. Plant Sci. 277: 155–165.

Nakayasu, M., Shioya, N., Shikata, M., Thagun, C., Abdelkareem, A., Okabe, Y., Ariizumi, T., Arimura, G.-i., Mizutani, M., Ezura, H., Hashimoto, T., and Shoji, T. (2018). JRE4 is a master transcriptional regulator of defense-related steroidal glycoalkaloids in tomato. Plant J. 94: 975–990.

Nakayasu, M., et al. (2020). Identification of α-tomatine 23-hydroxylase involved in the detoxification of a bitter glycoalkaloid. Plant Cell Physiol., in press (10.1093/pcp/pcz224).

Oberpichler, I., Rosen, R., Rasouly, A., Vugman, M., Ron, E.Z., and Lamparter, T. (2008). Light affects motility and infectivity of *Agrobacterium tumefaciens*. Environ. Microbiol. 10: 2020–2029.

Pauwels, L., De Clercq, R., Goossens, J., Iñigo, S., Williams, C., Ron, M., Britt, A., and Goossens, A. (2018). A dual sgRNA approach for functional genomics in Arabidopsis thaliana. G3: Genes, Genomes, Genet. 8: 2603–2615.

Ritter, A., et al. (2017). The transcriptional repressor complex FRS7-FRS12 regulates flowering time and growth in *Arabidopsis*. Nat. Commun. 8: 15235.

Ron, M., et al. (2014). Hairy root transformation using *Agrobacterium rhizogenes* as a tool for exploring cell type-specific gene expression and function using tomato as a model. Plant Physiol. 166: 455–469.

Sawai, S., et al. (2014). Sterol side chain reductase 2 is a key enzyme in the biosynthesis of cholesterol, the common precursor of toxic steroidal glycoalkaloids in potato. Plant Cell 26: 3763–3774.

Schweizer, F., Fernández-Calvo, P., Zander, M., Diez-Diaz, M., Fonseca, S., Glauser, G., Lewsey, M.G., Ecker, J.R., Solano, R., and Reymond, P. (2013). *Arabidopsis* basic helix-loop-helix transcription factors MYC2, MYC3, and MYC4 regulate glucosinolate biosynthesis, insect performance, and feeding behavior. Plant Cell 25: 3117–3132.

Sheard, L.B., et al. (2010). Jasmonate perception by inositol-phosphate-potentiated COI1–JAZ co-receptor. Nature 468: 400–405.

Shoji, T. (2019). The recruitment model of metabolic evolution: jasmonate-responsive transcription factors and a conceptual model for the evolution of metabolic pathways. Front. Plant Sci. 10: 560.

Shoji, T., and Hashimoto, T. (2011). Tobacco MYC2 regulates jasmonate-inducible nicotine biosynthesis genes directly and by way of the *NIC2*-locus *ERF* genes. Plant Cell Physiol. 52: 1117–1130.

Shoji, T., Ogawa, T., and Hashimoto, T. (2008). Jasmonate-induced nicotine formation in tobacco is mediated by tobacco *COI1* and *JAZ* genes. Plant Cell Physiol. 49: 1003–1012.

Shoji, T., Kajikawa, M., and Hashimoto, T. (2010). Clustered transcription factor genes regulate nicotine biosynthesis in tobacco. Plant Cell 22: 3390–3409.

Sonawane, P.D., et al. (2018). Short-chain dehydrogenase/reductase governs steroidal specialized metabolites structural diversity and toxicity in the genus *Solanum*. Proc. Natl. Acad. Sci. USA 115: E5419–E5428.

Sonawane, P.D., et al. (2016). Plant cholesterol biosynthetic pathway overlaps with phytosterol metabolism. Nat. Plants 3: 16205.

Spyropoulou, E.A., Haring, M.A., and Schuurink, R.C. (2014). RNA sequencing on *Solanum lycopersicum* trichomes identifies transcription factors that activate terpene synthase promoters. BMC Genomics 15: 402.

Sun, J.-Q., Jiang, H.-L., and Li, C.-Y. (2011). Systemin/jasmonate-mediated systemic defense signaling in tomato. Mol. Plant 4: 607–615.

Swinnen, G., Jacobs, T., Pauwels, L., and Goossens, A. (2020). CRISPR-Cas-mediated gene knockout in tomato. Methods Mol. Biol. 2083: 321–341.

Thagun, C., et al. (2016). Jasmonate-responsive ERF transcription factors regulate steroidal glycoalkaloid biosynthesis in tomato. Plant Cell Physiol. 57: 961–975.

Thines, B., Katsir, L., Melotto, M., Niu, Y., Mandaokar, A., Liu, G., Nomura, K., He, S.Y., Howe, G.A., and Browse, J. (2007). JAZ repressor proteins are targets of the SCF^COI1^ complex during jasmonate signalling. Nature 448: 661–665.

Townsley, B.T., Covington, M.F., Ichihashi, Y., Zumstein, K., and Sinha, N.R. (2015). BrAD-seq: Breath Adapter Directional sequencing: a streamlined, ultra-simple and fast library preparation protocol for strand specific mRNA library construction. Front. Plant Sci. 6: 366.

Van Bel, M., Diels, T., Vancaester, E., Kreft, L., Botzki, A., Van de Peer, Y., Coppens, F., and Vandepoele, K. (2018). PLAZA 4.0: an integrative resource for functional, evolutionary and comparative plant genomics. Nucleic Acids Res. 46: D1190–D1196.

Vanden Bossche, R., Demedts, B., Vanderhaeghen, R., and Goossens, A. (2013). Transient expression assays in tobacco protoplasts. Methods Mol. Biol. 1011: 227–239.

Vanholme, R., et al. (2013). Caffeoyl shikimate esterase (CSE) is an enzyme in the lignin biosynthetic pathway in *Arabidopsis*. Science 341: 1103–1106.

Wasternack, C., and Strnad, M. (2019). Jasmonates are signals in the biosynthesis of secondary metabolites — Pathways, transcription factors and applied aspects — A brief review. New Biotechnol. 48: 1–11.

Xu, J., van Herwijnen, Z.O., Dräger, D.B., Sui, C., Haring, M.A., and Schuurink, R.C. (2018). SlMYC1 regulates type VI glandular trichome formation and terpene biosynthesis in tomato glandular cells. Plant Cell 30: 2988–3005.

Xu, S., et al. (2017). Wild tobacco genomes reveal the evolution of nicotine biosynthesis. Proc. Natl. Acad. Sci. USA 114: 6133–6138.

Yamanaka, T., Vincken, J.-P., Zuilhof, H., Legger, A., Takada, N., and Gruppen, H. (2009). C22 Isomerization in α-tomatine-to-esculeoside a conversion during tomato ripening is driven by C27 hydroxylation of triterpenoidal skeleton. J. Agric. Food Chem. 57: 3786–3791.

Yan, J., Yao, R., Chen, L., Li, S., Gu, M., Nan, F., and Xie, D. (2018). Dynamic perception of jasmonates by the F-box protein COI1. Mol. Plant 11: 1237–1247.

Yan, J., Zhang, C., Gu, M., Bai, Z., Zhang, W., Qi, T., Cheng, Z., Peng, W., Luo, H., Nan, F., Wang, Z., and Xie, D. (2009). The *Arabidopsis* CORONATINE INSENSITIVE1 protein is a jasmonate receptor. Plant Cell 21: 2220–2236.

Zouine, M., Maza, E., Djari, A., Lauvernier, M., Frasse, P., Smouni, A., Pirrello, J., and Bouzayen, M. (2017). TomExpress, a unified tomato RNA-Seq platform for visualization of expression data, clustering and correlation networks. Plant J. 92: 727–735.

